# TRPV4 modulates substrate stiffness mechanosensing and transcellular pore formation in human Schlemm’s canal cells

**DOI:** 10.1101/2025.05.25.656000

**Authors:** Haiyan Li, Cydney Wong, Seyed Mohammad Siadat, Kristin M. Perkumas, Jacques A. Bertrand, Darryl R. Overby, Todd Sulchek, W. Daniel Stamer, C. Ross Ethier

**Affiliations:** Biomedical Engineering, Georgia Institute of Technology/Emory University, Atlanta, GA, United States; Department of Ophthalmology, Emory University, Atlanta, GA, United States; Department of Ophthalmology, Duke University, Durham, NC, United States; Bioengineering, Imperial College London, London, United Kingdom; Mechanical Engineering, Georgia Institute of Technology, Atlanta, GA, United States

**Keywords:** Glaucoma, mechanobiology, ion channel, calcium signaling, tissue engineering, hydrogel, ECM stiffening, trancellular pore

## Abstract

Pathological changes in the biomechanical environment of Schlemm’s canal (SC) inner wall cells, such as substrate stiffening and increased cellular stretch, are associated with ocular hypertension, a key risk factor for the development of glaucoma. Cell membrane stretch can trigger the activation of transient receptor potential vanilloid 4 (TRPV4) mechanosensitive ion channels, allowing calcium influx and initiating downstream signaling. However, the precise role of TRPV4 in SC cell mechanobiology remains unclear. Here, we demonstrate that sustained inhibition of TRPV4 activity modulates substrate stiffness mechanosensing to thereby affect the remodeling of the actin cytoskeleton and extracellular matrix of SC cells. This is accompanied by a reduction in cell stiffness and an increase in transcellular pore forming ability, potentially lowing outflow resistance and risk of ocular hypertension. Conversely, acute activation of TRPV4 channels induces Ca^2+^ influx, increasing transcellular pore formation in SC cells. Notaly, reduced TRPV4 mechanosensing was observed in glaucomatous SC cells, resulting in reduced transcellular pore forming ability. These findings suggest novel potential strategies based on targeting TRPV4 in SC cells for the treatment of ocular hypertension in glaucoma.

## Introduction

Glaucoma is the leading cause of irreversible vision loss worldwide, and elevated intraocular pressure (IOP) is the only modifiable risk factor for glaucoma^1–3^. IOP is largely determined by the resistance to aqueous humor (AH) outflow in the juxtacanalicular region of the conventional outflow pathway where the trabecular meshwork and Schlemm’s canal (SC) inner wall cells interact^4^. Dysfunction in the outflow tract impairs the eye’s ability to maintain IOP homeostasis, resulting in increased resistance to AH outflow and ocular hypertension (OHT).

SC is a continuous vessel lined with endothelial cells having phenotypic characteristics of both vascular and lymphatic endothelia^5–9^. Like lymphatics, cells forming the inner wall of SC experience a basal-to-apical pressure gradient that deforms these cells to create giant vacuoles and form pores that facilitate fluid drainage from the eye^10^. In primary open-angle glaucoma (POAG), the most common type of glaucoma, pathological changes in SC associated with OHT have been observed, including: a smaller lumen^11^, stiffer SC inner wall cells^12,13^, and decreased inner wall pore size and density^12,14,15^. However, the mechanisms responsible for differences in responses to mechanical stimuli between normal versus glaucomatous SC inner wall cells remain unclear.

The physiological substrate of SC inner wall endothelial cells is the outermost layer of the trabecular meshwork, which is rich in extracellular matrix (ECM) and is delimited by a discontinuous basement membrane underlying the inner wall. In eyes with POAG, the trabecular meshwork demonstrates elevated contractility and ECM deposition, leading to tissue stiffening^13,16–20^. Consequently, SC inner wall cells in the diseased outflow tract are subject to increased biomechanical stimuli from the stiffened trabecular meshwork. In addition, these cells are continuously subjected to IOP-induced biomechanical stresses (e.g., tension, fluid shear), leading to stretching of their plasma membranes, and these effects are greater when IOP is elevated. Notably, previous studies demonstrate that SC cells in culture alter the actin component of their cytoskeleton, ECM production, and cellular stiffness in response to a glaucomatous biophysical microenvironment^12,21^. It is believed that these cell-driven alterations lead to decreased outflow and consequently elevated IOP. Jointly, these pathological mechanisms can push the outflow tract to exceed its adaptive homeostatic capacity in a feed-forward loop.

Tissue health and disease are frequently governed by a complex interplay of cell-intrinsic and systemic factors, encompassing both biophysical and biochemical cues. The cellular membrane establishes a physical boundary between intracellular components and the extracellular environment, serving as a crucial interface that orchestrates responses to mechanical cues. Within cellular membranes, there resides a diverse array of mechanosensitive proteins, including integrins and mechanosensitive channels such as the transient receptor potential vanilloid 4 (TRPV4)^22–24^. TRPV4 channels are activated by mechanical stimuli such as pressure, tension, shear stress, changes in osmolarity, substrate stiffness, and pharmacological activators to allow calcium influx and thus initiate downstream signaling^24–28^. TRPV4 channels are involved in several physiological functions, such as regulating blood flow, shear stress-induced vasodilation, angiogenesis, epidermal permeability, tissue fibrosis development, and neuronal excitability^25^. Further, TRPV4 channel activity has been shown to affect functions of various ocular tissues, including the corneal epithelium, lens, ciliary body, retina, and trabecular meshwork^29–35^. Additionally, several *in vitro* and *in vivo* studies suggest that TRPV4 may play a role in glaucoma by modulating AH outflow and IOP^32,36,37^, particularly through its effects on trabecular meshwork cells, or by influencing retinal ganglion cell survival^38,39^. However, the precise role of TRPV4 in SC cells remains unclear. IOP affects the stretch of SC inner wall cells, which is expected to influence TRPV4 activity and cell function. Here we mimic IOP-induced TRPV4 activity modulation by using small molecule agonist/antagonist to explore the role of TRPV4 in regulating the function of healthy and glaucomatous human SC inner wall cells. In this manner, we investigated the potential role of TRPV4 in SC inner wall cells as a matrix stiffness sensor, influencing SC cell mechanobiology.

## Results

### TRPV4 activity affects SC cell mechanobiology

To determine the potential role(s) of TRPV4 in SC cell mechanobiology, we first confirmed its expression in primary normal human SC (nSC) cells derived from healthy donors (**Suppl. Fig. 1A**). We then tested whether TRPV4 activation would induce Ca^2+^ influx by using the specific agonist GSK101. GSK101 increased peak Ca^2+^ levels and caused sustained Ca^2+^ elevations (measured at 250 seconds post-GSK101 application) in nSC cells. This intracellular influx was largely abolished when SC cells were co-treated with GSK101 and the TRPV4 antagonist HC06. We also observed that HC06 treatment alone had no detectable effect on intracellular Ca^2+^ levels in nSC cells within the measured timeframe (**Suppl. Fig. 1B**).

TRPV4 has been shown to affect actin remodeling within the trabecular meshwork^32^. To determine whether a sustained reduction in TRPV4 activity also affects the actin network in SC cells, we treated nSC cells with the TPRV4 antagonist HC06. Following a 4-day treatment with HC06, we observed a significant reduction in F-actin stress fibers (44.5%, p<0.0001) in nSC cells (**Figure 1A, B**) and decreased αSMA expression at both the transcript and protein levels compared to vehicle controls (**Figure 1A, C; Suppl. 2A**). Upon removal of HC06 for 2 days, partial reversal of HC06-induced actin cytoskeletal remodeling was observed. Additionally, subsequent activation of TRPV4 by the agonist GSK101 for 2 days further enhanced actin remodeling, although it did not reach control levels (**Figure 1A-C; Suppl. 2A**).

**Fig. 1.**
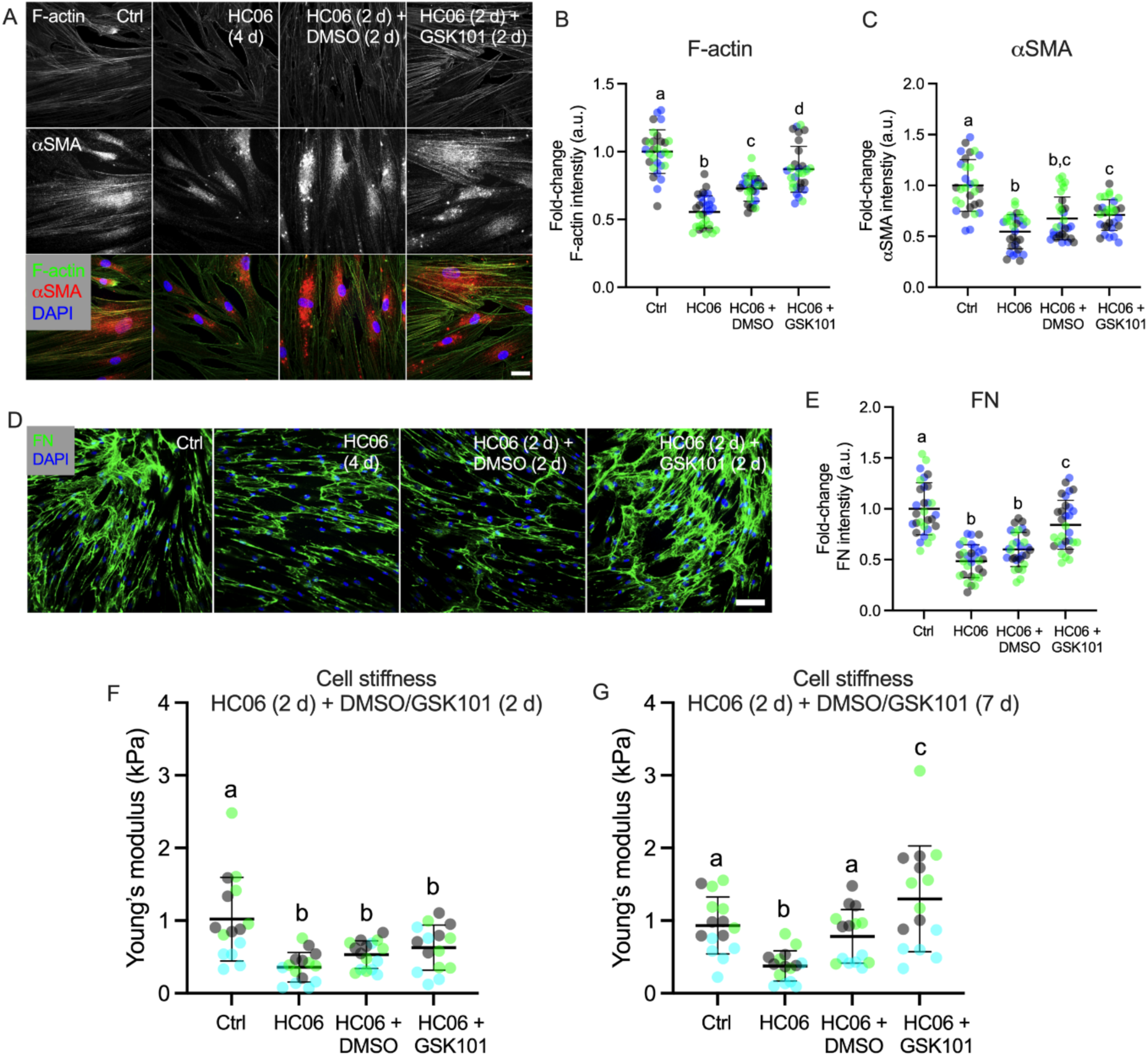
TRPV4-mediated mechanobiological response in normal Schlemm’s canal (nSC) cells. **(A)** Representative fluorescence micrographs of F-actin and αSMA in nSC cells under the following conditions: control (DMSO); exposure to the TRPV4 antagonist HC06 (10 μM) for 4 days; exposure to the TRPV4 antagonist HC06 (10 μM) for 2 days, followed by removal of HC06 and exposure to vehicle (DMSO) for 2 days; and exposure to the TRPV4 inhibitor HC06 (10 μM) for 2 days followed by removal of HC06 and exposure to the TRPV4 agonist GSK101 (100 nM) for 2 days. Scale bar, 20 μm. **(B, C)** Plots of F-actin and αSMA normalized protein labelling intensity (n = 30 images per group from 3 nSC cell strains with three replicates per cell strain). **(D)** Representative fluorescence micrographs showing fibronectin (FN) expression and deposition by nSC cells. Scale bar, 100 μm. **(E)** Analysis of FN protein labelling intensity (n = 30 images per group from 3 nSC cell strains with three replicates per cell strain). **(F)** Young’s modulus of nSC cells measured by AFM, with groups as in panel A (n = 15 cells per group from 3 nSC cell strains). **(G)** Young’s modulus of nSC cells with groups as in panel A, except that exposure to DMSO or GSK101 was for 7 days instead of 2 days (n = 15 cells per group from 3 nSC cell strains). Symbols with the same colors are from the same cell strains. Lines and error bars indicate mean ± SD. Significance was determined by two-way ANOVA using multiple comparisons tests [shared significance indicator letters represent non-significant difference (p > 0.05), distinct letters represent significant difference (p < 0.05)].

We next considered the impact of TRPV4 activity on ECM produced by SC cells. As was the case for actin, modulation of TRPV4 activity affected the amount of the SC inner wall basement membrane protein fibronectin, at both mRNA and protein levels (**Figure 1D, E**). Collagen types I and IV have been previously detected in the basement membrane of the inner wall of SC^40^, and we demonstrated that their transcript levels were also affected by TRPV4 activity in nSC cells (**Suppl. Fig. 2C, D**). Importantly, we also observed that TRPV4 activity influenced nSC cell stiffness, with inhibition of TRPV4 activity softening nSC cells, whereas activation of TRPV4 channels for 2 days after removing the inhibitor showed no significant effect on cell stiffness (**Figure 1F**).

We further investigated whether a longer duration of TRPV4 activation could fully reverse the effects induced by a 2-day inhibition of TRPV4 activity. Therefore, after inhibiting TRPV4 activity for 2 days with the antagonist HC06 as above, HC06 was withdrawn from the culture media and nSC cells were allowed to recover in control media or were treated with the TRPV4 agonist GSK101 for 7 days. In the recovery experiment, we observed that actin remodeling and the production/deposition of ECM proteins (i.e., fibronectin, collagen types I and IV) returned nearly to control levels after removing HC06 for 7 days. Additionally, TRPV4 activation for 7 days after 2 days of TRPV4 inhibition significantly increased actin and ECM remodeling above baseline levels (**Suppl. Fig. 3**). Importantly, similar trends were observed for cell stiffness, with 7 days of TRPV4 channel activation significantly increasing cell stiffness compared to the group in which the TRPV4 antagonist was removed (1.30 ± 0.73 kPa vs. 0.78 ± 0.37 kPa; **Figure 1G**). Taken together, these results suggest that long-term activation of TRPV4 channels influences SC cell stiffness by modifying actin levels and influences ECM remodeling.

### TRPV4 modulates nSC cell substrate stiffness mechanosensing

Previous studies have indicated that the normal trabecular meshwork has a stiffness of approximately 0.5-9 kPa, as measured by atomic force microscopy^13,16–18^ (**Suppl. Table 1**). In contrast, the trabecular meshwork in glaucoma eyes is ∼1.5-20-fold stiffer, depending on the measurement methods used and the location of measurement^13,16–18^. As a result, SC inner wall cells in the diseased outflow tract experience an altered microenvironment due to their underlying stiffened substrate. To study the interactions between substrate stiffness and TRPV4 channels in SC cells, we grew cells on fibronectin-coated, tissue-mimetic hydrogel substrates that were either soft or stiff (elastic moduli of 2.36 kPa and 8.00 kPa, respectively, as measured by AFM).

Consistent with previous studies^12,41^, we observed that nSC cells on the stiff hydrogel substrate exhibited increased F-actin stress fibers and cell stiffness compared to cells on the soft substrate (**Suppl. Fig. 4**). Intracellular calcium concentration is known to have an important role in the organization of F-actin^42,43^. Thus, we examined levels of intracellular calcium in nSC cells on both soft and stiff substrates, finding that steady-state intracellular Ca^2+^ concentration was significantly higher in cells residing on the stiff substrate vs. in cells on the soft substrate (1.33-fold, p<0.001; **Figure 2A, B**). Additionally, nSC cells residing on the stiff substrates exhibited elevated TRPV4 transcript levels vs. in cells on the soft substrate (1.31-fold, p<0.01; **Suppl. Figure 4D**). This suggests that increased TRPV4 levels on stiff substrates may, in part, contribute to increased intracellular Ca^2+^ levels.

**Fig. 2.**
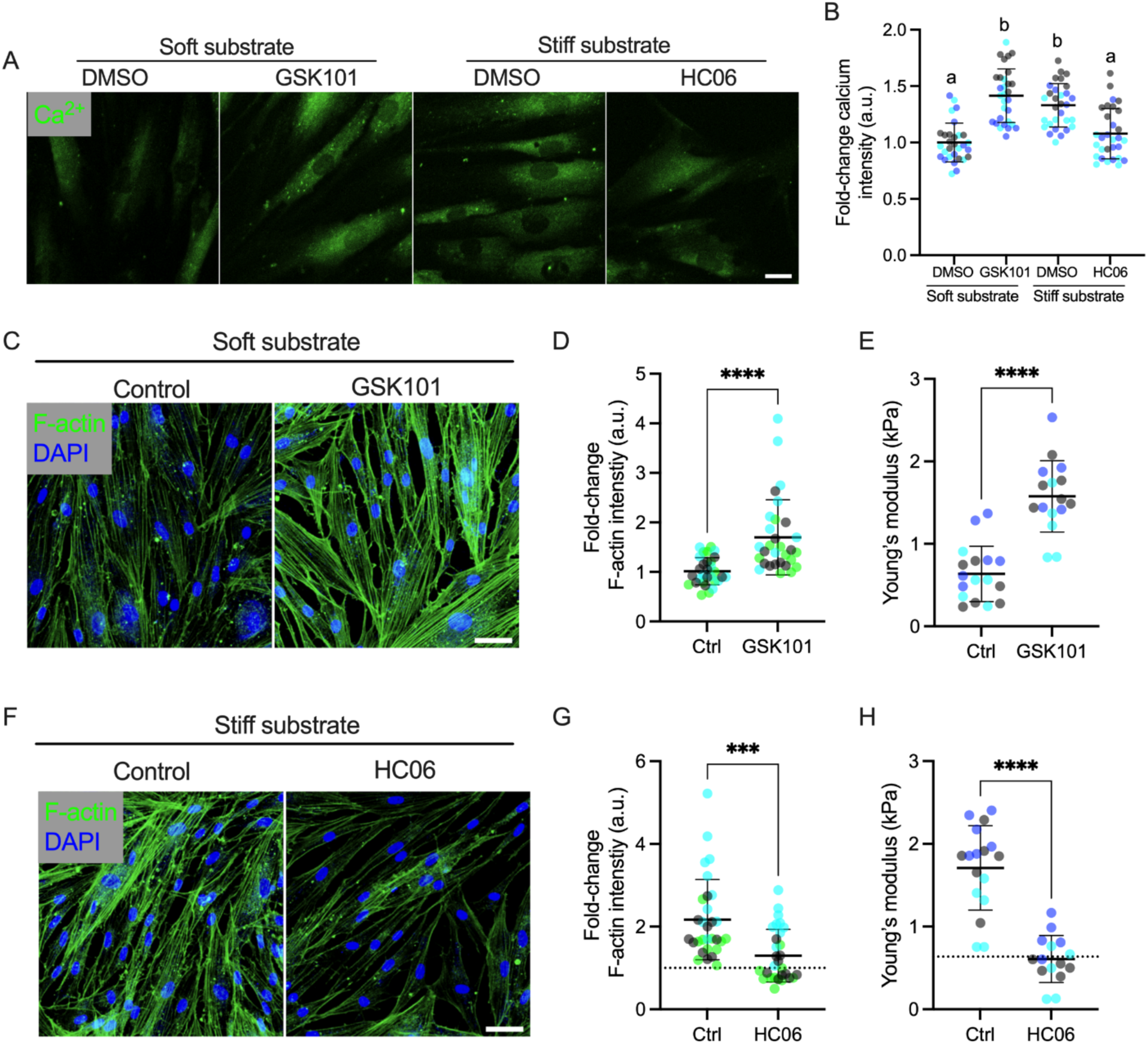
TRPV4 activity in nSC cells affects their response to substrate stiffness. **(A)** Representative fluorescence micrographs of intracellular Ca^2+^ in Fluo-4AM-labeled nSC cells on the soft and stiff hydrogel substrates under control (DMSO) conditions, or treated with the TRPV4 agonist GSK101 (100 nM) or antagonist HC06 (10 μM) for 2 days. Scale bars, 20 μm. **(B)** Plots of normalized intracellular Ca^2+^ fluorescence intensity (n = 10 images per group from 3 nSC cell strains). **(C)** Representative fluorescence micrographs of F-actin in nSC cells on the soft substrate grown for 2 days in either control (DMSO) conditions, or with the TRPV4 agonist GSK101 (100 nM) added to the media. Scale bar, 20 μm. **(D, E)** Plots of F-actin fluorescent intensity and Young’s modulus measured by AFM in nSC cells exposed to control media or the TRPV4 agonist GSK101 for 2 days (100 nM; F-actin: n = 30 images per group from 3 nSC cell strains with three replicates per cell strain; AFM: n = 16-17 cells per group from 3 nSC cell strains). **(F)** Representative fluorescence micrographs of F-actin in nSC cells grown on the stiff substrate for 2 days in either control (DMSO) conditions, or with the TRPV4 antagonist HC06 (10 μM) added to the media. Scale bar, 20 μm. **(G, H)** Similar to panels (D, E), except that cell were grown on the stiff substrate and treated with a TRPV4 antagonist for 2 days. Symbols with the same colors are from the same cell strains. The lines and error bars indicate mean ± SD; dotted lines show the control mean values obtained from nSC cell grown on the soft substrate, for reference. Significance was determined by two-way ANOVA using multiple comparisons tests (***p<0.001, ****p<0.0001).

To further investigate the role of TRPV4 channels in mechanosensing of substrate stiffness by nSC cells, we treated cells residing on the soft substrate with the TRPV4 agonist GSK101, followed by assessing intracellular Ca^2+^, F-actin, and cell stiffness. In the converse experiment, we also treated nSC cells residing on the stiff substrate with the TRPV4 antagonist HC06. We observed that sustained TRPV4 activation in nSC cells residing on soft substrates significantly increased intracellular Ca^2+^ levels, leading to calcium levels comparable to those seen in cells grown on the stiff substrate (**Figure 2A, B**). Conversely, sustained TRPV4 inhibition in cells residing on the stiff substrate decreased Ca^2+^ to levels comparable to those seen in cells residing on the soft substrate (**Figure 2A, B**). Additionally, TRPV4 activation increased F-actin stress fibers in nSC cells grown on the soft substrate (1.85-fold, p<0.0001; **Figure 2C, D**) and induced cell stiffening (p<0.0001; **Figure 2E**), while TRPV4 inhibition in cells grown on the stiff substrate reduced F-actin levels (0.60-fold, p<0.0001; **Figure 2E,F**) and cell stiffness (p<0.0001; **Figure 2G**), bringing them to levels comparable to those observed in nSC cells on the soft hydrogels. Taken together, these observations show that decreased levels of intracellular Ca^2+^ caused by sustained TRPV4 inhibition shift nSC cells grown on the stiff substrate toward the phenotype observed on the soft substrate, and vice versa. These findings are consistent with the concept that TRPV4 may play a role in modulating substrate stiffness responses in nSC cells.

### Glaucomatous SC cells show impaired responses to TRPV4 modulation

Previous studies have shown that glaucomatous SC (gSC) cells (derived from patients with glaucoma) are stiffer and exhibit a more fibrotic phenotype compared to cells from non-glaucomatous donors^12,44^. Here, we observed that gSC cells exhibited significantly lower TRPV4 transcript levels than nSC cells (**Figure 3A**), and decreased levels of peak and sustained intracellular Ca^2+^ in response to activation by GSK101 (**Figure 3B**).

**Fig. 3.**
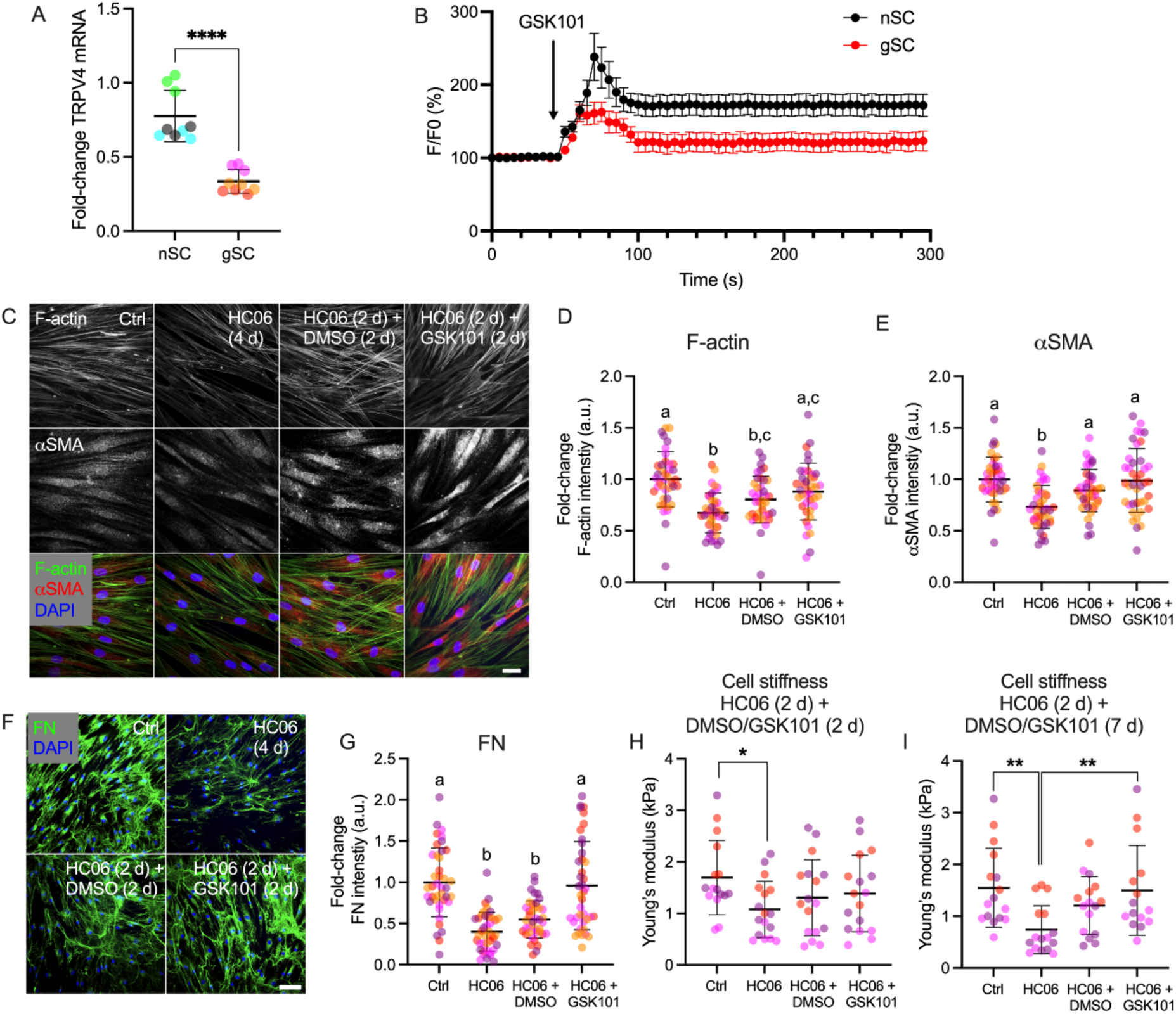
Responses of glaucomatous SC (gSC) cells toTRPV4 modulation. **(A)** Normalized TRPV4 mRNA levels in normal (nSC) and glaucomatous (gSC) SC cells, determined by qRT-PCR (normalization to GAPDH levels; n = 9 replicates from 3 nSC/3 gSC cell strains). **(B)** Relative fluorescence intensities (F/F0) of the calcium indicator Fluo-4AM in nSC and gSC cells in response to the TRPV4 agonist GSK101 (100 nM; n = 12 replicates from 3 nSC cell strains, n = 4 replicates from 2 gSC cell strains). **(C)** Representative fluorescence micrographs of F-actin and αSMA in gSC cells under the following conditions: control (DMSO); exposure to the TRPV4 antagonist HC06 (10 μM) for 4 days; exposure to the TRPV4 antagonist HC06 (10 μM) for 2 days, followed by removal of HC06 and exposure to vehicle (DMSO) for 2 days; and exposure to the TRPV4 inhibitor HC06 (10 μM) for 2 days followed by removal of HC06 and exposure to the TRPV4 agonist GSK101 (100 nM) for 2 days. Scale bar, 20 μm. **(D, E)** Plots of F-actin and αSMA fluorescent labeling intensities (n = 40 images per group from 4 gSC cell strains with three replicates per cell strain). **(F)** Representative fluorescence micrographs of fibronectin (FN) in gSC cells cultured under various conditions as in panel A. Scale bar, 100 μm. **(G)** Plots of FN fluorescence intensity (n = 40 images per group from 4 gSC cell strains with three replicates per cell strain). **(H, I)** Young’s modulus of gSC cells measured by AFM with either 2 (panel H; n = 16-17 cells per group from gSC cell strains) or 7 days treatment with TRPV4 agonist (panel I; n = 15-16 cells per group from 3 gSC cell strains). Symbols with the same colors are from the same cell strains. The lines and error bars indicate mean ± SD. Significance was determined by two-way ANOVA using multiple comparisons tests [*p<0.05, **p<0.01, shared significance indicator letters represent non-significant difference (p > 0.05), distinct letters represent significant difference (p < 0.05)].

We next asked how gSC cells respond to sustained modulation of TRPV4 activity. Similar to our observations in nSC cells, a 4-day treatment with the TRPV4 antagonist HC06 significantly decreased actin levels in gSC cells, although the magnitude of the change was less than that seen in nSC cells. Removal of the TRPV4 antagonist HC06 after 2 days restored actin to vehicle control levels. Two days of TRPV4 channel activation with GSK101 after 2 days of inhibition with HC06 showed no significant effect on F-actin, and 7 days of activation with GSK101 significantly increased F-actin levels compared to the group with HC06 removal and DMSO addition (**Figure 3C-E; Suppl. Fig. 5, Suppl. Fig. 6A-C**).

Interestingly, 4 days of TRPV4 channel inhibition did not affect fibronectin expression in gSC cells at the transcript level (**Suppl. Fig. 5A**), but there was a significant decrease in fibronectin expression/deposition at the protein level (**Fig. 3F, G**), suggesting perhaps that TRPV4 may influence proteins associated with fibronectin degradation. No significant change in fibronectin was detected with HC06 removal, and both 2 days and 7 days of TRPV4 channel activation significantly increased fibronectin at the protein level, but not at the mRNA level (**Fig. 3F, G; Suppl. Fig. 5, Suppl. Fig. 6D, E**). We also observed that TRPV4 activation in gSC cells increased the mRNA levels of collagen types I and IV, but these effects were dampened compared to those seen in nSC cells (**Suppl. Fig. 5**).

Consistent with a previous report^12^, gSC cells were measured to be stiffer than nSC cells by AFM (**Figure 3H, I**). Comparable to the trends observed with F-actin, both TRPV4 inhibition and activation modulated cell stiffness to a lesser extent in gSC cells vs. in nSC cells. Specifically, 4 days of TRPV4 inhibition decreased nSC cell stiffness by 64.7%, while it reduced gSC cell stiffness by only 36.6%; and 7 days of TRPV4 activation increased cell stiffness by 1.58-fold in nSC cells, but only by 1.25-fold in gSC cells (**Figure 1F, G, Figure 3H, I**). Overall, these results demonstrate that, in contrast to normal SC cells, gSC cells exhibit an impaired response to TRPV4 modulation.

Next, we sought to investigate the interplay between TRPV4 activity and substrate stiffness mechanosensing in gSC cells. Similar to observations in nSC cells, gSC cells exhibited higher levels of F-actin and cell stiffness on the stiff substrate compared to the soft substrate (**Figure 4**). Unexpectedly, sustained TRPV4 activation significantly increased F-actin, but not stiffness, in gSC cells on the soft substrate (**Figure 4A-C**), while sustained TRPV4 inhibition showed no effect on either F-actin or stiffness in gSC cells on the stiff substrate (**Figure D-F**). Taken together, these results confirm that interactions between TRPV4 activity and the response to substrate stiffness are muted in glaucomatous SC cells as compared to normal SC cells.

**Fig. 4.**
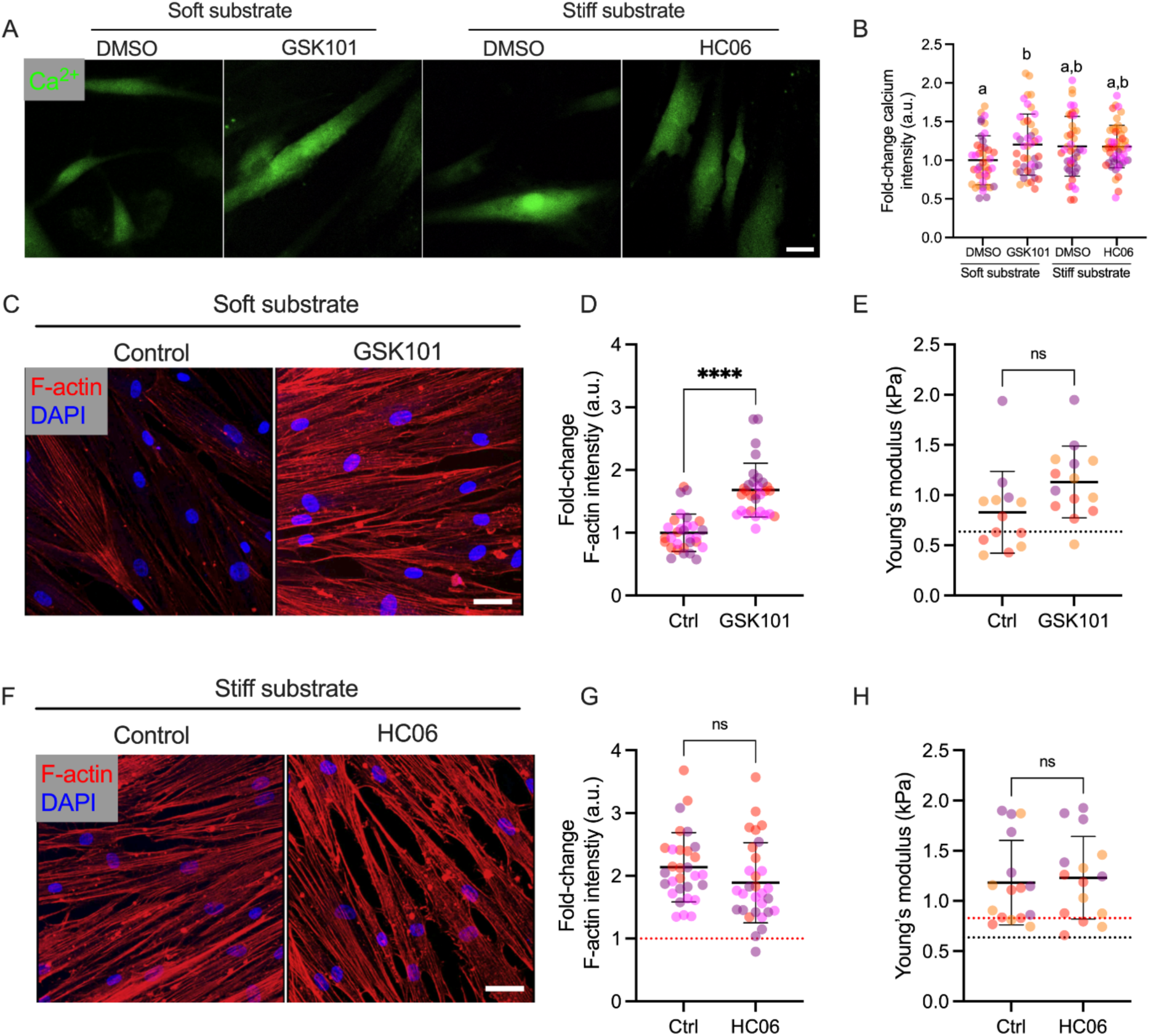
Impaired TRPV4-modulated substrate stiffness mechanosensing in gSC cells. **(A)** Representative fluorescence imaging of intracellular Ca^2+^ in Fluo-4AM-labeled gSC cells cultured on the soft and stiff hydrogel substrates, under control conditions or after modulation of TRPV4 activity for 2 days. Scale bars, 20 μm. **(B)** Plots of normalized intracellular Ca^2+^ fluorescence intensity corresponding to conditions shown in panel A (n = 40 images per group from 4 gSC cell strains with three replicates per cell strain). **(C)** Representative fluorescence micrographs of F-actin in gSC cells on the soft substrate grown for 2 days in either control (DMSO) conditions, or with the TRPV4 agonist GSK101 (100 nM) added to the media. Scale bar, 20 μm. **(D, E)** Plots of F-actin fluorescent intensity and Young’s modulus measured by AFM in gSC cells exposed to control media or the TRPV4 agonist GSK101 for 2 days (100 nM; F-actin: n = 30 images per group from 3 gSC cell strains with three replicates per cell strain; AFM: n = 16-17 cells per group from 3 gSC cell strains). **(F)** Representative fluorescence micrographs of F-actin in gSC cells grown on the stiff substrate for 2 days in either control (DMSO) conditions, or with TRPV4 antagonist HC06 (10 μM) added to the media. Scale bar, 20 μm. **(G, H)** Plots of F-actin fluorescent intensity and Young’s modulus measured by AFM in gSC cells exposed to control media or the TRPV4 antagonist HC06 for 2 days (10 μM; F-actin: n = 30 images per group from 3 gSC cell strains with three replicates per cell strain; AFM: n = 15-17 cells per group from 3 gSC cell strains). Symbols with different colors represent different cell strains. The lines and error bars indicate Mean ± SD; black and red dotted lines show nSC and gSC cell on the soft substrate control mean value for reference. Significance was determined by two-way ANOVA using multiple comparisons tests (****p<0.0001).

### Acute TRPV4 activation increases transcellular pore formation

To drain from the eye via SC, aqueous humor passes through micron-sized pores in the inner wall of SC; further, these pores play a crucial role in modulating IOP^7,12,45^. Thus, there is great interest in understanding mechanobiological signaling processes that modulate SC cell pore formation. It is also well known that IOP fluctuations (on time scales of seconds to days^46,47^) cause stretching of the plasma membranes of SC inner wall cells, which in turn is expected to activate TRPV4 mechanosensitive ion channels and promote Ca^2+^ influx. We hypothesized that acute TRPV4 activation-induced intracellular Ca^2+^ elevation could influence pore formation in SC cells, thereby modulating outflow facility and IOP. Membrane fusion is required for transcellular pore formation, and emerging evidence suggests that intracellular Ca^2+^ dynamics is a key modulator of membrane fusion and fusion-pore expansion^48^.

To assess transcellular pore formation in SC cells, our lab has previously developed an *in vitro* assay that subjects SC cells to localized cellular stretching^49^. In this assay, carboxyl microspheres (4.0-4.9 μm diameter) are seeded on a biotinylated gelatin substrate, followed by plating of SC cells. Subsequently, fluorescently-tagged streptavidin is introduced into the culture medium. The fluorescently-tagged streptavidin traverses the SC cell monolayer at locations where pores (or gaps between cells) exist, thereby irreversibly adhering to the biotinylated substrate and thus generating an observable persistent fluorescent signal (**Figure 5A**).

**Fig. 5.**
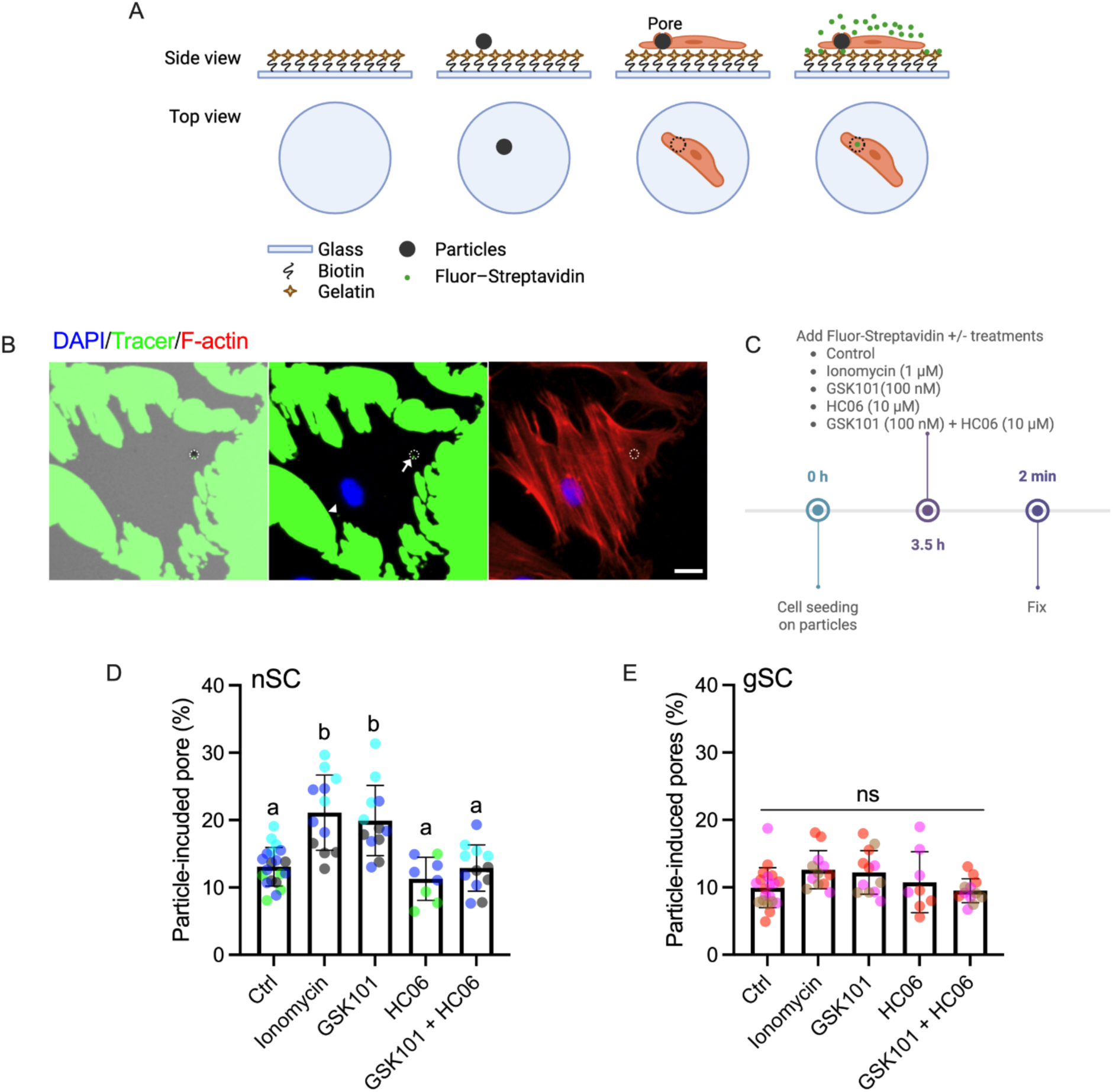
Rapid TRPV4 activation induces upregulation of intracellular Ca^2+^ and enhances trancellular pore formation in SC cells. **(A)** Schematic showing particles for inducing, and fluorescence assay for detecting, pores in SC cells. Carboxyl particles (4.0-4.9 μm diameter) were seeded on a biotinylated gelatin substrate, followed by plating of normal or glaucomatous SC cells. Subsequently, fluorescently-tagged streptavidin was introduced into the culture media. **(B)** Representative fluorescent micrographs showing spots where fluorescently tagged streptavidin (green) has bound with biotin substrate under cultured SC cells (at sites of intracellular pores) as well as surrounding regions not covered by cells. White circles outline particles. The arrow indicates a particle-induced pore, while the arrowhead points to a spontaneously formed pore detected by the fluorescent assay but not associated with a particle. Scale bar: 50 μm. **(C)** Schematic showing the time course of pore formation assay: SC cells were seeded atop particles on a biotinylated gelatin-coated glass substrate. Three and half hours after cell seeding, fluorescently-tagged streptavidin plus the relevant treatment (DMSO [control], 1 μM ionomycin, 100 nM GSK101, 10 μM HC06, or co-treatment with 10 μM HC06 plus 100 nM GSK101) was introduced into the culture media and incubated for 2 mins. Created with BioRender.com. **(D)** Pore incidence (percentage of particles associated with pores) in nSC cells (n = 4 wells per nSC cell strain). **(E)** Pore incidence in gSC cells (n = 4 wells per gSC cell strain). Symbols with different colors represent different cell strains. The lines and error bars indicate Mean ± SD. Significance was determined by two-way ANOVA using multiple comparisons tests [shared significance indicator letters represent non-significant difference (p > 0.05), distinct letters represent significant difference (p < 0.05)].

Using our pore forming assay, we observed that basal-to-apical cellular stretching imparted by particles underlying SC cells caused transcellular pore formation (**Figure 5B**). To investigate the acute effects of TRPV4 activation on transcellular pore formation in SC cells, GSK101 was added to the media overlying cells along with the fluorescent tracer. Ionomycin, an effective Ca^2+^ ionophore, was used as a positive control for intracellular Ca^2+^ induction (**Figure 5C**). In normal cells, 13.1% of particles exhibited an associated pore, and both ionomycin and TRPV4 activation significantly increased particle-induced pore formation, while co-treatment of HC06 with GSK101 blocked TRPV4 activation-induced trancellular pore formation. Notably, HC06 treatment alone showed no effect on SC cell transcellular pore formation, aligning with the observation that HC06 exhibited no impact on intracellular Ca^2+^ levels during short-term treatment. These data are consistent with the hypothesis that TRPV4 activation can promote transcellular pore formation in normal SC cells by upregulating intracellular Ca^2+^ (**Figure 5D**).

In glaucomatous SC cells, 9.9% of particles were associated with pores, which was significantly lower than the frequency observed in normal cells, consistent with previously observed differences in trancellular pore formation between normal vs. glaucomatous SC cells, both in native tissues and in an *in vitro* assay inducing trancellular pore formation by fluid flow across SC cell monolayers ^12,14^. A similar pattern of TRPV4 activation and ionomycin effects on trancellular pore formation was observed in gSC cells as in nSC cells (**Figure 5E**). Consistent with our previous observations, the effects of TRPV4 activity modulation/intracellular Ca^2+^ were attenuated in gSC cells compared to normal cells. Specifically, in nSC cells, ionomycin and GSK101 increased trancellular pore formation rate by 1.61- and 1.52-fold compared to control conditions, respectively, whereas these increases were only 1.27- and 1.23-fold in gSC cells. Together, these results indicate that acute TRPV4 activation-induced upregulation of intracellular Ca^2+^ enhances trancellular pore formation in SC cells, with greater efficiency observed in normal cells compared to glaucomatous cells.

### Extended TRPV4 inhibition promotes trancellular pore formation in nSC cells, but not in gSC cells

Previous studies have shown a strong inverse association between trancellular pore formation and cell stiffness in SC cells^12,50^. In this study, we have demonstrated that long-term inhibition of TRPV4 activity decreases SC cell stiffness. Therefore, we hypothesized that long-term inhibition of TRPV4 channels would also elevate the ability of SC cells to form transcellular pores. To test this hypothesis, SC cells were treated with HC06 in high serum conditions for two days. We used high serum for these experiments, since we observed that low serum conditions affected cell spreading time, which in turn adversely impacted the pore detection assay. After treatment, the cells were trypsinized and plated on particles to induce pore formation using the assay described above (**Figure 6A**).

**Fig.6.**
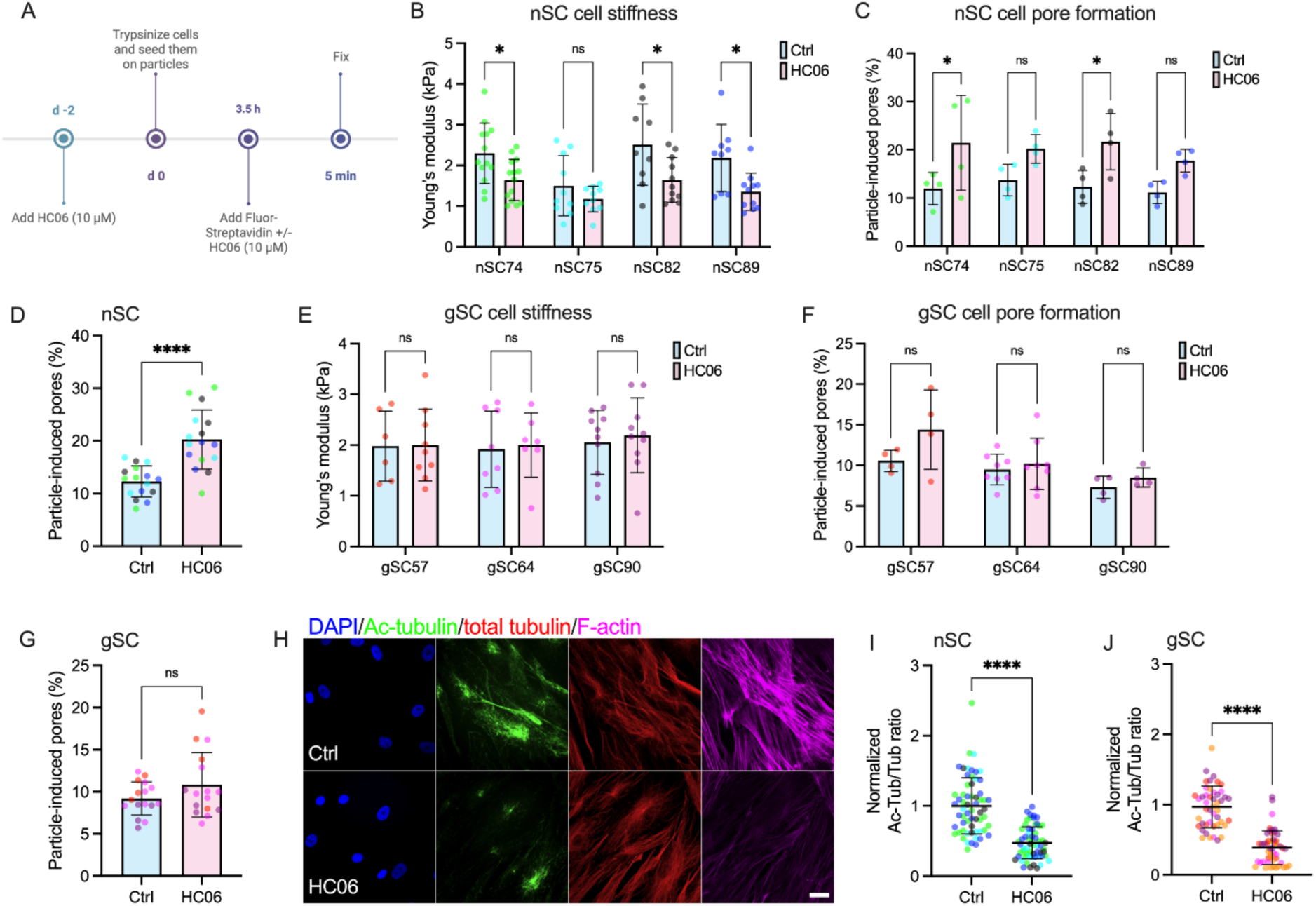
Impact of prolonged TRPV4 inhibition on stiffness, trancellular pore formation, and MTs in SC cells. **(A)** Schematic illustrating the pore formation assay timeline: SC cells were treated with 10 μM HC06 in medium containing 10% FBS for 2 days, followed by trypsinization and plating on particles to induce pore formation. Cells were allowed to spread for 3.5 hours under either control (DMSO) conditions, or with the TRPV4 antagonist HC06 (10 μM) added to the media, and transcellular pore formation was detected using fluorescently-tagged streptavidin. Created with BioRender.com. **(B, E)** Young’s modulus of nSC and gSC cells measured by AFM (nSC cell: n = 9-14 cells per group; gSC cell: n = 6-10 cells per group). **(C, F)** Pore formation rate (percentage of particles with an associated pore) induced by particles in nSC (n = 4 wells per group) and gSC cells (n = 4-8 wells per group). **(D, G)** Pooled data on transcellular pore formation rate from all nSC and gSC cell strains. Symbols with different colors represent different cell strains. **(H)** Representative fluorescence micrographs of F-actin, acetylated α-tubulin (Ac-tubulin), and total α-tubulin in nSC cells treated with DMSO or TRPV4 antagonist HC06 (10 μM). Scale bar, 20 μm. **(I, J)** Ratio of acetylated to total microtubule fluorescent intensity in normal and gaucomatous SC cells (n = 20 images per group from 2 nSC/gSC cell strains with three replicates per cell strain). Symbols with different colors represent different cell strains. The lines and error bars indicate Mean ± SD. Significance was determined by two-way ANOVA using multiple comparisons tests (*p<0.05, ****p<0.0001).

First, we confirmed that actin and fibronectin levels remained depressed by TRPV4 inhibition after trypsinization in both nSC and gSC cells (**Suppl. Fig. 7**). By measuring cell stiffness using AFM, and by measuring trancellular pore formation rate as described above, we also observed that inhibiting TRPV4 for two days in high serum conditions significantly decreased cell stiffness in three out of the four normal cell strains tested (**Figure 6B**). The exception was nSC75 cells, in which cell stiffness was decreased from 1.50 ± 0.74 kPa to 1.18 ± 0.32 kPa, but this difference did not reach statistical significance. Using our pore forming assay, we found that trancellular pore formation rate was increased in two of the four cell strains tested, with the other strains showing a trend that did not reach statistical significance (**Figure 6C**). However, despite the variability in particle-induced pore formation rates across the tested cell strains, pooled data showed that long-term TRPV4 inhibition significantly increased particle-induced pore formation in nSC cells compared to vehicle controls (20.3 ± 5.6% vs. 12.3 ± 2.9%; **Figure 6D**).

We next repeated these experiments in gSC cells. Interestingly, no significant changes were observed in either cell stiffness or particle-induced pore formation for any of the individual gSC cell strains tested (**Figure 6E, F**). Pooling the data from all cell strains revealed a small increase in trancellular pore formation rate associated with TRPV4 inhibition (10.8 ± 3.8% vs. 9.2 ± 2.0%); however, this difference was not statistically significant (**Figure 6G**).

Sustained TRPV4 inhibition downregulated F-actin in both normal and glaucomatous cells, but this did not translate to significant changes in cell stiffness. We thus investigated whether other cytoskeletal elements that could influence SC cell stiffness, specifically microtubules (MTs)^51^, were modulated by TRPV4 activity. MT stability has been shown to influence actin organization, cell contractility, and stiffness in other cell types^52–57^. Further, acetylation is a proxy for MT stability^58^; therefore, we assessed the effects of TRPV4 inhibition on MT acetylation. By analyzing the ratio of acetylated α-tubulin to total α-tubulin, we found that sustained TRPV4 inhibition significantly decreased MT stability in both normal and glaucomatous SC cells (**Figure 6H-J**). This suggests that changes in MT stability, along with F-actin changes, may influence cell stiffness. Overall, these data suggest that long-term TRPV4 inhibition increases the rate of particle-induced pore formation in nSC cells by modulating cell stiffness, but it is not effective in gSC cells.

## Discussion

In this study, we show that TRPV4 activity modulates SC cell mechanobiology, including F-actin architecture, ECM protein expression/deposition, cell stiffness, and transcellular pore formation. In particular, we observe that human SC cells sense their substrate stiffness, and that this interaction is modulated at least in part by TRPV4 (**Figure 7**). Our data support the notion that acute TRPV4 activation induces Ca^2+^ influx, increasing transcellular pore formation in SC cells to facilitate outflow and decrease IOP in normal SC cells. Paradoxically, sustained TRPV4 inhibition in normal SC cells also increases transcellular pore formation. The situation in glaucomatous SC cells is slightly different: there are lower transcript levels of TRPV4, and the effects of TRPV4 activation and inhibition are attenuated as compared to the situation in normal SC cells.

**Figure 7.**
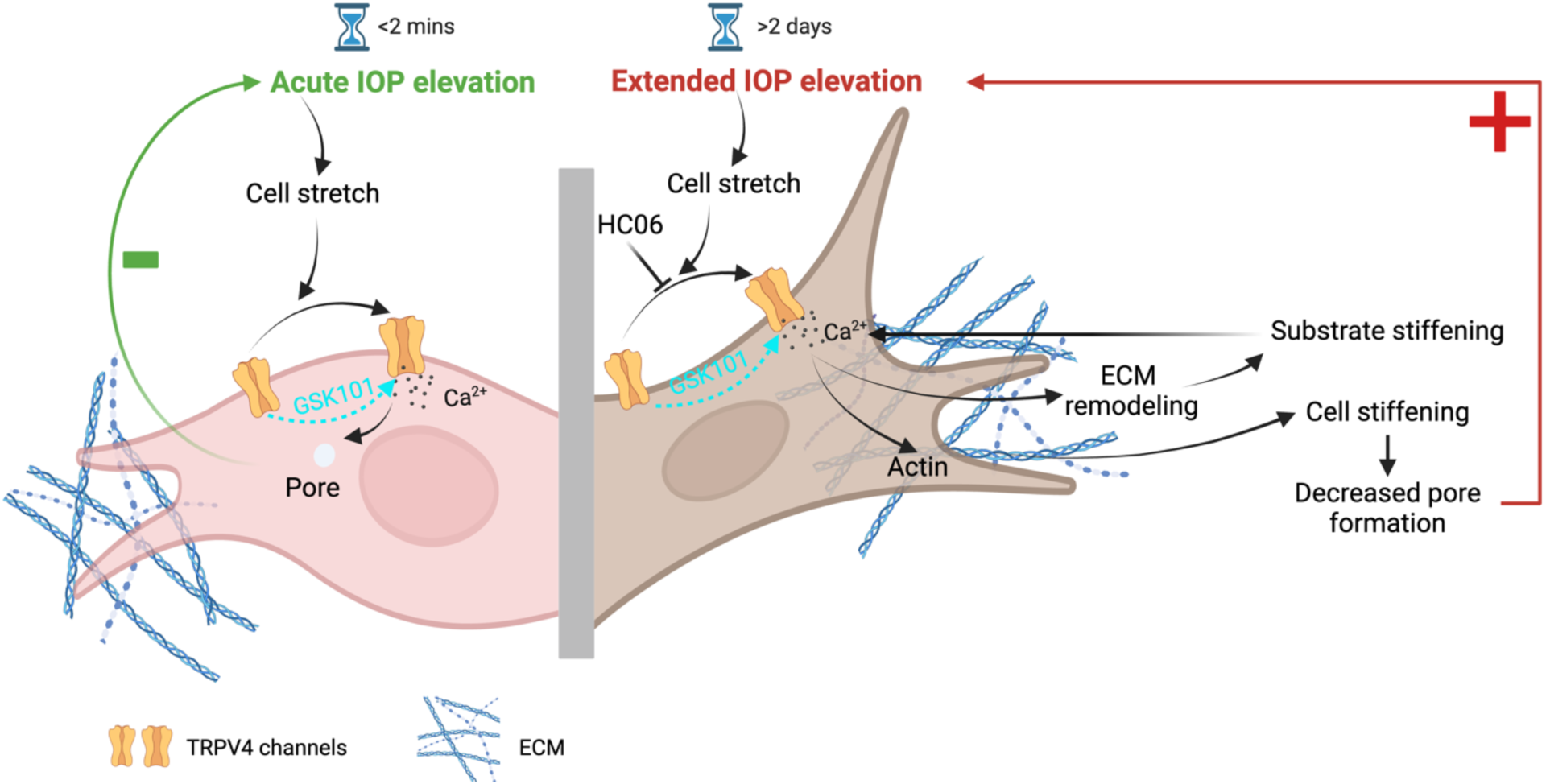
Schematic for hypothesized role of TRPV4 mechanotransduction in SC cells. Under normal circumstances (left), acute IOP elevation activates TRPV4 channels, inducing Ca^2+^ influx and transcellular pore formation to facilitate aqueous outflow, thereby decreasing IOP and maintaining homeostasis. However, if IOP is chronically elevated (right), the SC inner wall cell substrate stiffens which induces elevation of intracellular Ca^2+^, this leads to actin and ECM remodeling, increasing cell stiffness and forming a positive feedback loop that mitigates against IOP homeostasis. Sustained TRPV4 modulation influences SC cell mechanobiology, where inhibition reduces stiffness and increases pore formation, while activation has the opposite effect. The effects of TRPV4 modualtion are diminished in glaucomatous SC cells. Created with BioRender.com.

In glaucoma, we postulate that impaired TRPV4 function in SC cells makes cells unable to counteract acute IOP transients, leading to sustained IOP elevation. Long-term IOP elevation induces stiffening of the SC inner wall cell substrate and TRPV4 activation, resulting in SC cell cytoskeletal and ECM remodeling. This causes SC cell stiffening, in turn reducing their pore formation ability, and creates a positive feedback loop that further increases IOP (**Figure 7**).

A recent study identified an association between human genetic risk variants in mechanosensitive channels, such as TRPV4, and glaucoma^59^. Additionally, TRPV4 channels have been shown to influence IOP and conventional outflow facility in both mice^60^ and *in vitro* studies, where trabecular meshwork cells were cultured on porous SU-8 polymers, by modulating trabecular meshwork cell function^32^. Our data demonstrated that TRPV4 activity also influences the mechanobiology of human SC cells in the outflow tract. Specifically, sustained inhibition of TRPV4 activity decreased F-actin and αSMA levels and the expression/deposition of ECM proteins (e.g., fibronectin, collagen types I and IV), which could be reversed by prolonged withdrawal of the TRPV4 inhibitor or by subsequent TRPV4 activation. αSMA, fibronectin, and collagen type I are markers of endothelial-to-mesenchymal transition (EndMT)^61–63^, suggesting that sustained TRPV4 inhibition has the potential to block the EndMT of SC cells, thereby inhibiting cell pathology.

The SC inner wall experiences a basal-to-apical pressure gradient which deforms SC cells to create giant vacuoles and transcellular pores that facilitate fluid drainage from the eye^10^. Our lab has previously developed an *in vitro* platform that utilizes micron-sized particles to mimic the giant vacuoles and to apply basal-to-apical local cellular stretch, demonstrating the associated formation of transcellular pores in SC cells. Importantly, using this assay, we observed that the ability of SC cells to form pores significantly associated with cell stiffness^50^. In this study, we extended these findings to determine that prolonged TRPV4 inhibition led to SC cell softening and increased transcellular pore formation in SC cells. These data suggest that long-term TRPV4 inhibition has the potential to increase outflow facility. Therefore, the reduced IOP in mouse eyes induced by a TRPV4 antagonist could be due to both trabecular meshwork relaxation and increased SC endothelial permeability.

Changes in the mechanical properties of the SC inner wall substrate have been implicated in the initiation and progression of glaucoma. For example, the substrate of the SC inner wall, known as the trabecular meshwork, is markedly stiffer in glaucoma^13,16,18^. To model this situation, we fabricated biomimetic hydrogel systems with stiffnesses mimicking the situation in normal and glaucomatous eyes. We observed that SC cells exhibited a higher level of intracellular Ca^2+^, F-actin fibers, and cell stiffness on stiff substrates vs. on soft substrates, consistent with previous studies^41^. These studies thus identify TRPV4 channels as a modulator, and perhaps even a direct mechanosensor, of substrate stiffness sensing in SC cells. Further, on soft substrates, sustained TRPV4 activation increased SC cell stiffness, while conversely, TRPV4 inhibition reversed the cell stiffening induced by stiff substrates. While the data clearly implicate TRPV4 as a modulator of calcium levels and cell phenotype in response to matrix stiffness, other glaucoma-related mechanosensitive channels, such as PIEZO, TRPV2, and TRPM3^59^, may also play a role in the mechanotransduction in SC cells.

IOP is known to fluctuate over time scales ranging from seconds to days^46,47^. Increased IOP causes stretching of the SC inner wall cell plasma membranes and thus very likely activates TRPV4. Previous studies have reported that acute cellular stretch (i.e., 2 minutes) induces pore formation in SC cells, which can be blocked by depleting endoplasmic reticulum or extracellular calcium^64,65^. Our data revealed that acute TRPV4 activation increased particle-induced pore formation in SC cells, which was blocked by co-treatment of TRPV4 antagonist with an agonist. Interestingly, we observed that treatment with the TRPV4 antagonist alone had no significant effect on transcellular pore formation. Several factors may explain this observation: (i) the experimental sequence, in which mechanical stretch from particles was applied before antagonist treatment, may have activated TRPV4 channels and triggered Ca^2+^ influx prior to inhibition; thus the antagonist may not have reversed early Ca^2+^ signaling events within the short experimental timeframe; (ii) compensatory activity from other mechanosensitive ion channels, such as PIEZO channels, could mask the effect of TRPV4 inhibition; and (iii) mechanical stretch may have induced internalization of TRPV4 channels^66^, reducing their membrane localization and thereby limiting the efficacy of the antagonist. Topical administration of a TRPV4 agonist has been shown to lower nocturnal IOP in mouse eyes within 30 minutes of application^60^. We thus suggest that the rapid decrease in IOP following TRPV4 activation may be attributed to enhanced transcellular pore formation in the inner wall of SC.

Interestingly, our studies revealed that both TRPV4 activation and inhibition effects in SC cells derived from glaucomatous patients were diminished compared to observations in normal cells, as evidenced by smaller/no changes in cell stiffness and pore formation due to perturbation in TRPV4 activity. This reduction could be due to the decreased TRPV4 message levels and activity we observed in gSC cells. TRPV4 is known to internalize in response to agonist stimulation^66^. In glaucoma, persistently elevated IOP may lead to chronic TRPV4 activation, triggering its internalization and thereby reducing its functional activity. Our observations indicate that TRPV4 activation induces stiffening in normal SC cells. In contrast, gSC cells are inherently stiffer and exhibit lower TRPV4 activity compared to normal cells, suggesting additional factors influencing cell stiffness are altered in SC cells under glaucomatous conditions. These observations highlight the complexity of SC cell mechanobiology in glaucoma, and therapeutic strategies targeting TRPV4 may need to be tailored to address these specific alterations.

In conclusion, we demonstrate treating normal SC cells cultured on stiff substrates with a TRPV4 antagonist significantly decreased cell stiffness and elevated transcellular pore formation ability, thereby protecting normal cells from acquiring a glaucomatous SC-like phenotype. These results are in agreement with findings that TRPV4 knockout protects mouse eyes from developing high IOP^32^. Collectively, our data reveal that while TRPV4 has a key role in mechanotransduction in normal SC cells, this function seems to be impaired in glaucomatous SC cells. It is crucial to determine the timing and mechanism of the molecular changes in glaucomatous SC cells that impairs their response to TRPV4 activity modulation. Such insights will inform efforts to restore this function in glaucomatous SC cells and thus may guide future interventions in glaucoma pathology.

## Methods

### SC cell isolation and culture

Primary human SC cells were isolated from ostensibly normal and glaucomatous human donor eyes, as previously described, and were cultured according to established protocols^9,63^. A total of five normal SC cell (nSC) strains isolated from healthy donor eyes and four glaucomatous cell (gSC) strains isolated from donors with a history of glaucoma were used in this study (**Table 1**). All SC cell strains were characterized based upon their typical spindle-like elongated cell morphology, expression of vascular endothelial-cadherin and fibulin-2, a net transendothelial electrical resistance of 10 ohms·cm^2^ or greater, and lack of myocilin induction following exposure to dexamethasone. Different combinations of SC cell strains were used per experiment depending on cell availability, and all studies were conducted between cell passage 3-6. SC cells were cultured in low-glucose Dulbecco’s Modified Eagle’s Medium (DMEM; Gibco; Thermo Fisher Scientific) containing 10% fetal bovine serum (FBS; HyClone, Thermo Fisher Scientific) and 1% penicillin/streptomycin/glutamine (PSG; Gibco) and maintained at 37°C in a humidified atmosphere with 5% CO_2_. Fresh media was supplied every 2-3 days.

**Table 1.** SC cell strain information. NM = not measured. N/A = not applicable. The first letter of the ID indicates normal (“n”) or glaucomatous (“g”)

| ID | Age | Sex | Facility<br>($\mu\text{L}/\text{min}/\text{mm Hg}$) | Ophthalmic<br>Medications |
| --- | --- | --- | --- | --- |
| nSC74 | 8 months | Male | NM | N/A |
| nSC75 | 10 years | Male | NM | N/A |
| nSC82 | 56 years | Male | NM | N/A |
| nSC87 | 62 years | Male | NM | N/A |
| nSC89 | 68 years | Male | NM | N/A |
| gSC57 | 78 years | Male | NM | Unknown |
| gSC64 | 78 years | Male | 0.07 | Unknown |
| gSC90 | 71 years | Female | 0.25 | Cosopt and<br>Latanoprost |
| gSC95 | 78 years | Male | 0.1 | N/A |

### Hydrogel preparation

To prepare the soft substrate, the hydrogel precursor gelatin methacryloyl (GelMA [6% w/v final concentration], Advanced BioMatrix, Carlsbad, CA, USA) was mixed with lithium phenyl-2,4,6-trimethylbenzoylphosphinate (LAP, 0.075% w/v final concentration) photoinitiator (Sigma-Aldrich, Saint Louis, MO). Stiff substrate also incorporated methacrylate-conjugated hyaluronic acid (MA-HA; [0.25% w/v final concentration], Advanced BioMatrix). Thirty microliters of the hydrogel solution were pipetted onto Surfasil-coated (Thermo Fisher Scientific) 18 × 18-mm square glass coverslips followed by placing 12-mm round silanized glass coverslips on top to facilitate even spreading of the polymer solution. Hydrogels were crosslinked by exposure to UV light (CL-3000 UV Crosslinker; Analytik Jena, Germany) at 1J/cm^2^. The hydrogel-adhered coverslips were removed with fine-tipped tweezers and placed hydrogel-side facing up in 24-well culture plates (Corning; Thermo Fisher Scientific). All the hydrogel substrates were coated with fibronectin (5 μg/cm^2^; Advanced BioMatrix) at room temperature for 1 hour.

### SC cell seeding and treatments

SC cells were seeded at 2 × 10^4^ cells/cm^2^ on premade hydrogel substrates or glass coverslips and cultured in DMEM with 10% FBS and 1% PSG for one or two days until ∼80-90% confluence was reached. Then, SC cell-seeded hydrogels were cultured in DMEM with 10% FBS and 1% PSG plus the relevant activator or inhibitor, depending on the treatment protocol.

For cells grown on coverslips, the treatment paradigms were as follows: (i) control (DMSO), exposure to the TRPV4 antagonist HC067047 (HC06; 10 μM, Sigma-Aldrich) for 4 or 9 days, exposure to HC06 for 2 days followed by its removal and continued culture in control conditions for 2 or 7 days, and (iv) exposure to HC06 for 2 days followed by the removal of HC06 and addition of the TRPV4 agonist GSK1016790A (GSK101; 100 nM, Sigma-Aldrich) for 2 or 7 days.

For cells grown on hydrogels, the treatment paradigms were: (i) grown on soft substrates and treated with the TRPV4 agonist GSK101 (100 nM) for 2 days, or (ii) grown on stiff substrates and treated with the TRPV4 antagonist HC06 (10 μM) for 2 days.

### Immunocytochemistry staining analysis

SC cells were fixed with 4% paraformaldehyde (PFA; Thermo Fisher Scientific) at room temperature for 20 mins, permeabilized with 0.5% Triton™ X-100 for 10 mins (Thermo Fisher Scientific), blocked with goat serum (BioGeneX, Fremont, CA, USA), and incubated with primary antibodies (anti-TRPV4, LS-C95181-500, 1:100, LSBio; anti-Fibronectin, ab45688, Abcam, 1:500; Cy3-anti-αSMA, C6198, Sigma-Aldrich, 1:400; anti-alpha Tubulin (acetyl K40), ab179484, Abcam, 1:1000; anti-alpha Tubulin, ab7291, Abcam, 1:1000) followed by incubation with fluorescent secondary antibody (Alexa Fluor® 488 anti-Rabbit, A27034, Invitrogen, 1:500; Alexa Fluor® 546 anti-Mouse, A11003, Invitrogen, 1:500). Nuclei were counterstained with 4′,6′-diamidino-2-phenylindole (4 μM, DAPI; Abcam, Waltham, MA, USA). Similarly, cells were stained with Phalloidin-iFluor 488 (Invitrogen; Thermo Fisher Scientific) or 594 (1:500, Cell signaling technology)/DAPI (Abcam) according to the manufacturer’s instructions. Glass coverslips/hydrogel substrates were mounted with ProLong™ Gold Antifade (Invitrogen) on Superfrost™ microscope slides (Fisher Scientific), and fluorescent images were acquired with a Leica DM6 B upright microscope system or with a Zeiss LSM 700 confocal microscope system.

All fluorescent image analyses were performed using FIJI software^67^ (National Institutes of Health (NIH), Bethesda, MD, USA). Fluorescence intensity of all F-actin, fibronectin (FN), or alpha smooth muscle actin (αSMA) fibers within the image was measured with image background subtraction. We then normalized the intensity by cell number and calculated fold-difference vs. control cells.

### Quantitative reverse transcription-polymerase chain reaction (qRT-PCR) analysis

Total RNA was extracted from SC cells using PureLink RNA Mini Kit (Invitrogen; Thermo Fisher Scientific). RNA concentration was determined with a NanoDrop spectrophotometer (Thermo Fisher Scientific). RNA was reverse transcribed using iScript™ cDNA Synthesis Kit (BioRad, Hercules, CA, USA). One hundred nanograms of cDNA were amplified in duplicates in each 40-cycle reaction using a CFX 96 Real Time PCR System (BioRad) with annealing temperature set at 60°C, Power SYBR™ Green PCR Master Mix (Thermo Fisher Scientific), and custom-designed qRT-PCR primers (Table 2). Transcript levels were normalized to GAPDH, and mRNA fold-change calculated relative to mean control values using the comparative CT method.

**Table 3.**
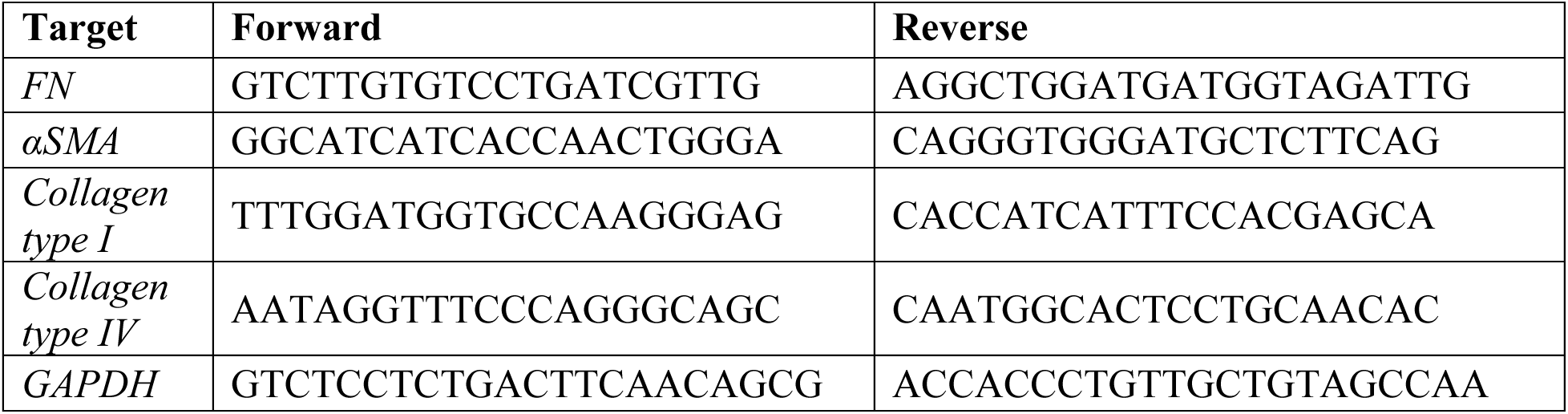
Oligonucleotide primer sequences (5’-3’) for qRT-PCR.

### Ca^2+^ imaging and analysis

SC cells were incubated with fluo-4AM (1 μM, Fisher Scientific) or fluo-520AM (5 μM, AAT Bioquest) in 1× HBSS solution (pH 7.4; Fisher Scientific) containing 10 mM HEPES (Fisher Scientific) for 45 minutes at 37°C in a humidified atmosphere with 5% CO_2_. This was followed by a 30-minute incubation with the staining buffer without the dye to remove any unloaded dye. The cells on hydrogels were then imaged using a Zeiss LSM 700 confocal microscope system.

For Ca^2+^ influx induction experiments, calcium influx was induced using the TRPV4 agonist (GSK101, 100 nM) or a combination of the TRPV4 agonist (GSK101, 100 nM) and antagonist (HC06, 10 μM). Images were captured every 5 seconds for 5 minutes, with the induction solution added after 90 seconds of imaging. Fluorescence intensity of intracellular Ca^2+^ was quantified using FIJI ImageJ. Cytosolic calcium increases (Ca^2+^ influx) were recorded by measuring ΔF/F (max-min), presented as relative fluorescence units.

### Cell and hydrogel stiffness measurement

SC cells were seeded at 2 × 10^4^ cells/cm^2^ on premade hydrogel substrates or glass coverslips and cultured in DMEM with 10% FBS and 1% PSG for one or two days until cells reached ∼80-90% confluence. Then, SC cells were exposed to various conditions depending on the experiment, as described above. An MFD-3D AFM (Asylum Research, Santa Barbara, CA, USA) was used to make stiffness measurements using silicon nitride cantilevers with an attached borosilicate sphere (diameter = 10 μm; nominal spring constant = 0.1 N/m; Novascan Technologies, Inc., Ames, IA, USA). Cantilevers were calibrated by measuring the thermally induced motion of the unloaded cantilever before measurements. The trigger force was set to 600 pN to avoid substrate effects and the tip velocity was adjusted to 800 nm/s to avoid viscous effects^68^. Five measurements/cell were conducted, and at least 5 cells were measured/group of one cell strain. For hydrogel stiffness measurement, a force map covering a 20 × 20 μm area (5 x 5 grid of points) was measured. Data from AFM measurements were fitted to the Hertz model to calculate the effective Young’s modulus of the cells, assuming the Poisson’s ratio was 0.5.

### Intracellular pore detection assay

16-well chambered glass bottom plates (Invitrogen) were plasma-treated and coated with 0.5 mg/mL biotinylated gelatin overnight at 37°C. The gelatin was then crosslinked with microbial transglutaminase (0.1 unit/µL, Sigma-Aldrich) at 37°C overnight. After UV sterilization of the plate, carboxyl ferromagnetic particles (4.0-4.9 µm; Spherotech Inc.) were added and incubated at room temperature for 20 minutes, followed by three gentle PBS washes. SC cells were seeded on top of the particles at a density of 7.5 × 10³ cells/cm² in DMEM supplemented with 10% FBS and 1% PSG, and incubated for 3.5 hours at 37°C with 5% CO_2_.

To acutely activate TRPV4 channels, cells were incubated with a fluorescent tracer (Streptavidin, Alexa Fluor™ 488 Conjugate; Invitrogen) under different treatments [DMSO, 1 μM ionomycin (Sigma-Aldrich), 100 nM GSK101, and a combination of 10 μM HC06 with 100 nM GSK101] in DMEM supplemented with 10% FBS and 1% PSG for 2 minutes. The cells were then immediately washed three times with PBS, fixed with 4% formaldehyde for 15 minutes at room temperature, permeabilized with 0.1% Triton X for 15 minutes, and stained with Phalloidin-594 (Cell Signaling Technology) and DAPI. The glass cover was detached from the 16-well chambered system using a coverglass removal tool (Invitrogen), mounted on another glass slide with Prolong Gold Antifade Mountant (Invitrogen), and stored in the dark at 4°C until examination with a Leica DM6 B upright microscope system. Fluorescent and brightfield images were acquired using a 10x objective to generate tile scans for each well.

For extended TRPV4 inhibition experiments, SC cells were pretreated with 10 μM HC06 in DMEM supplemented with 10% FBS and 1% PSG for 2 days, followed by trypsinization and plating on particles. After 3.5 hours of cell spreading, cells were incubated with the fluorescent tracer (Streptavidin, Alexa Fluor™ 488 Conjugate) under control (DMSO) or 10 μM HC06 conditions in DMEM supplemented with 10% FBS and 1% PSG for 5 minutes. The cells were then fixed, stained, and imaged as described above.

For the pore formation analysis, a 4 mm diameter circle was delineated at the center of each tile-scanned well image to exclude cells near the well edges, restricting the analysis to cells within this circle. The numbers of cells, particles beneath cells, and pores co-localized with particles were manually counted. The percentage of particle-induced pores was quantified as the number of pores associated with particles divided by the number of particles, multiplied by 100.

### Statistical analysis

Statistical analysis was conducted using GraphPad Prism, with significance set at p < 0.05. Sample sizes are specified in each figure caption. All data sets, except for AFM, were tested for normality using the Shapiro-Wilk test and were confirmed to meet the normality criteria. The AFM data sets, which are expected to be log-normally distributed^69^, were tested for log-normality using the Shapiro-Wilk test and met the relevant criteria. Comparisons between groups were assessed by one-way or two-way analysis of variance (ANOVA) with Tukey’s multiple comparisons *post hoc* tests, as appropriate.

## Disclosure

The authors report no conflicts of interest.

## Acknowledgments

This study was financially supported BrightFocus Foundation: CG2020001 (CRE), GR00028593 (HL); National Institutes of Health: 5R21EY033142-02 (CRE and DRO), R01EY028608 (WDS), R01EY022359 (WDS and DRO), T32 GM145735 (CAW), NIH Diversity Supplement EY031710-01S1 (CAW); BBSRC APP26196 (DRO); Georgia Research Alliance (CRE), and National Science Foundation Award Number 2134701 (TS) and CBET-2225476 (TS).

## Author contributions

**Conceptualization:** H.L., W.D.S., D.R.O., C.R.E.; **Investigation:**H.L., C.W., J.A.B., K.M.P., C.R.E. designed all experiments, collected, analyzed, and interpreted the data. C.W. assisted with AFM measurements and data analysis. J.A.B. helped with the pore formation assay. K.M.P. isolated and characterized the primary human SC cells; **Writing – original draft:** H.L., C.R.E.; **Writing – review and editing:** all authors. **Supervision:** C.R.E.

## Data and materials availability

All data needed to evaluate the conclusions in the paper are present in the paper and/or the Supplementary Materials. Additional data related to this paper may be requested from the authors.

## Supplementary Information

**Suppl. Fig. 1.**
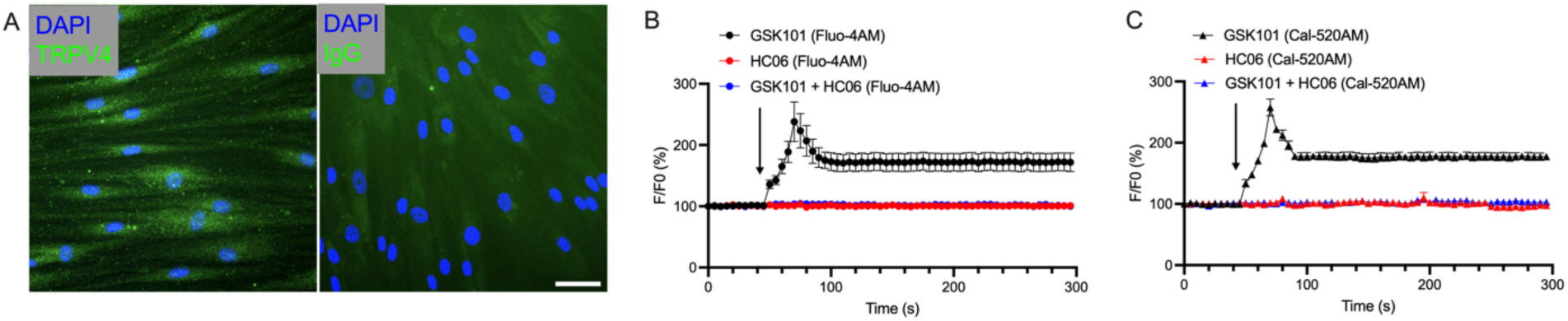
TRPV4 calcium channels are expressed and are functional in normal human SC cells. (A) Representative fluorescence micrographs of TRPV4 in nSC cells. The left image shows nSC cells stained with an anti-TRPV4 primary antibody, and the right image, serving as a negative control, shows cells stained with IgG. Nuclei are labeled blue. Scale bar, 50 μm. (B) The normalized fluorescence intensity (F/F0) of Fluo-4AM-loaded nSC cells is shown as a function of time in cells treated with the TRPV4 agonist GSK101 (100 nM; black line), TRPV4 antagonist HC06 (red line; 10 μM), or co-treated with the TRPV4 agonist GSK101 (100 nM) and the antagonist HC06 (blue line; 10 μM). The symbols and error bars indicate mean ± SEM (n = 12 replicates from 3 nSC cell strains). (C) The normalized fluorescence intensity (F/F0) of Cal-520AM-loaded nSC cells is shown as a function of time in cells. The symbols and error bars indicate mean ± SEM (n = 3 replicates from one nSC cell strain).

**Suppl. Fig. 2.**
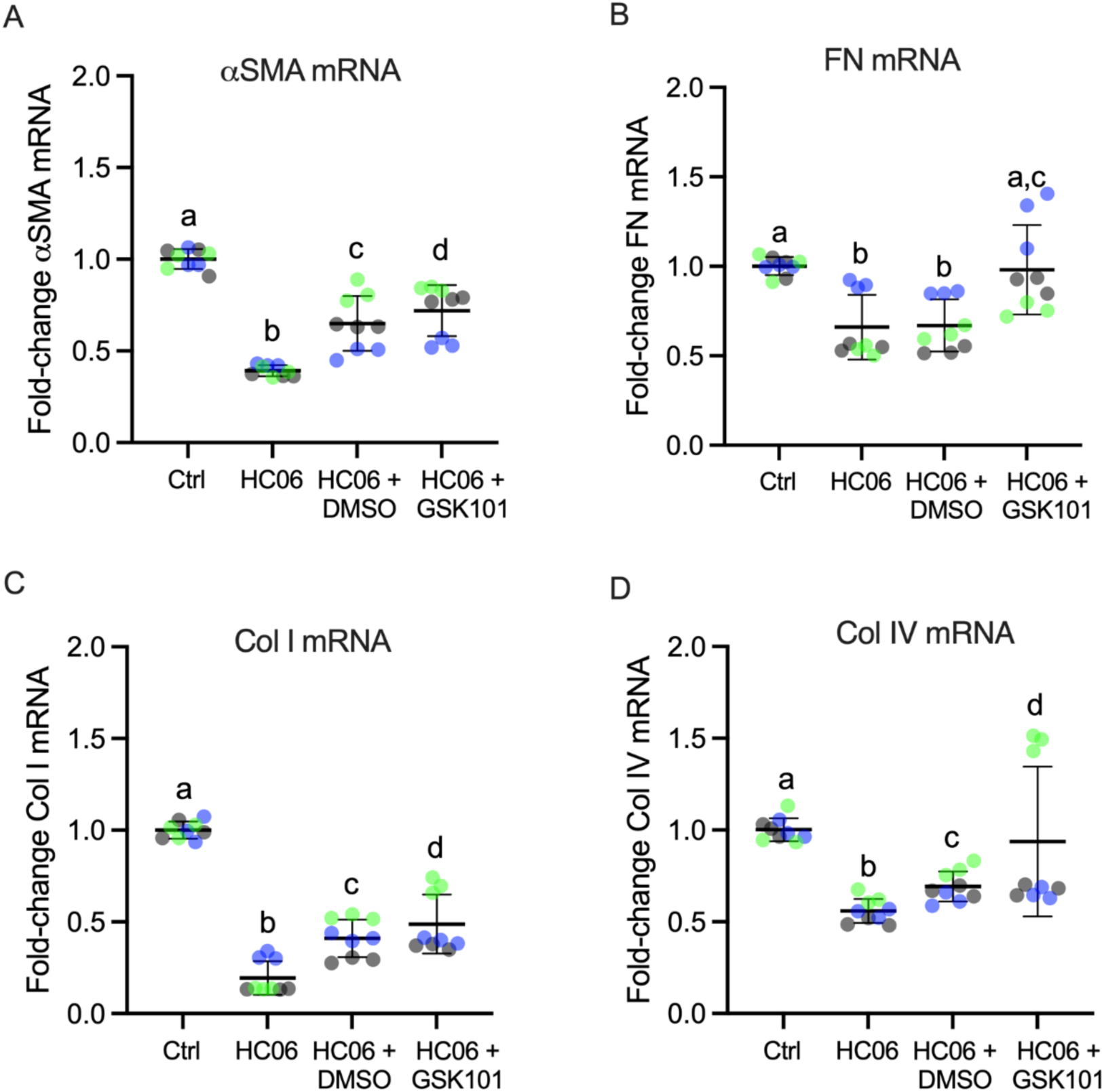
TRPV4 activity influences the transcript levels of αSMA and ECM proteins present in the basement membrane of the inner wall of SC. We plot the fold-change of mRNA levels for (**A**) αSMA, (**B**) fibronectin (FN), **(C)** collagen type I (Col I), and **(D)** collagen type IV (Col IV) normalized to GAPDH by qRT-PCR (n = 9 replicates from 3 nSC cell strains) in nSC cells under the following conditions: control (DMSO); exposure to the TRPV4 antagonist HC06 (10 μM) for 4 days; exposure to the TRPV4 antagonist HC06 (10 μM) for 2 days, followed by removal of HC06 and exposure to vehicle (DMSO) for 2 days; and exposure to the TRPV4 inhibitor HC06 (10 μM) for 2 days followed by removal of HC06 and exposure to the TRPV4 agonist GSK101 (100 nM) for 2 days. Symbols with the same colors are from the same cell strains. The lines and error bars indicate mean ± SD. Significance was determined by two-way ANOVA using multiple comparisons tests [shared significance indicator letters represent non-significant difference (p > 0.05), distinct letters represent significant difference (p < 0.05)].

**Suppl. Fig. 3.**
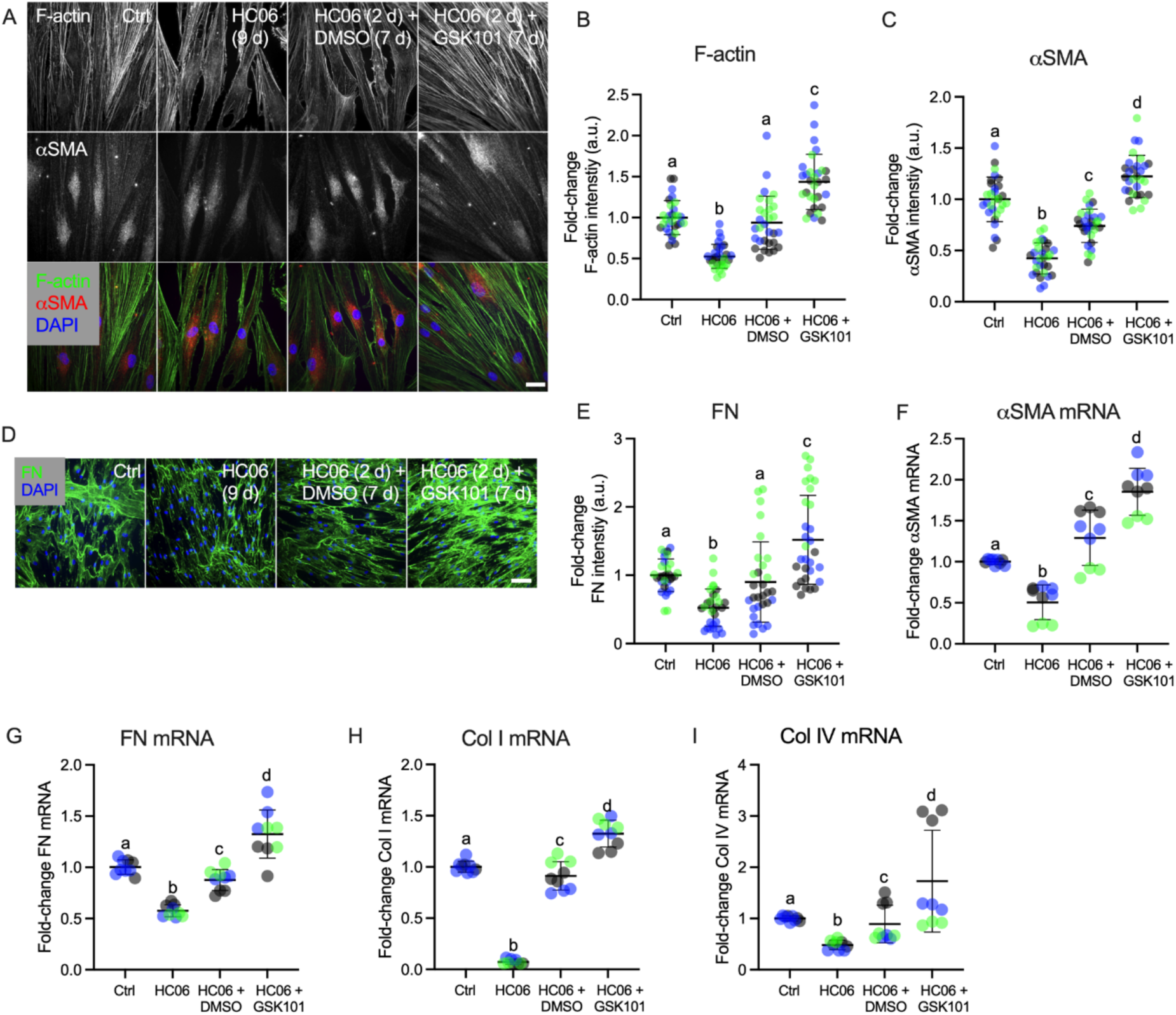
Extended TRPV4 channel activation reverses the mechanobiological changes in nSC cells induced by TRPV4 inhibition. **(A)** Representative fluorescence micrographs of F-actin and αSMA in nSC cells under the following conditions: control (DMSO); exposure to the TRPV4 antagonist HC06 (10 μM) for 9 days; exposure to the TRPV4 antagonist HC06 (10 μM) for 2 days, followed by removal of HC06 and exposure to vehicle (DMSO) for 7 days; and exposure to the TRPV4 inhibitor HC06 (10 μM) for 2 days followed by removal of HC06 and exposure to the TRPV4 agonist GSK101 (100 nM) for 7 days. Scale bar, 20 μm. **(B, C)** Plots of normalized F-actin and αSMA labelled protein intensity (n = 30 images per group from 3 nSC cell strains with three replicates per cell strain). **(D)** Representative fluorescence micrographs of fibronectin (FN) in nSC cells. Scale bar, 100 μm. **(E)** Plot of normalized FN protein labeling intensity (n = 30 images per group from 3 nSC cell strains with three replicates per cell strain). **(F-I)** Fold-change of mRNA levels for αSMA, fibronectin (FN), collagen type I (Col I), and collagen type IV (Col IV) normalized to GAPDH by qRT-PCR (n = 9 replicates from 3 nSC cell strains) in nSC cells. Symbols with the same colors are from the same cell strains. The lines and error bars indicate mean ± SD. Significance was determined by two-way ANOVA using multiple comparisons tests [shared significance indicator letters represent non-significant difference (p > 0.05), distinct letters represent significant difference (p < 0.05)].

**Suppl. Fig. 4.**
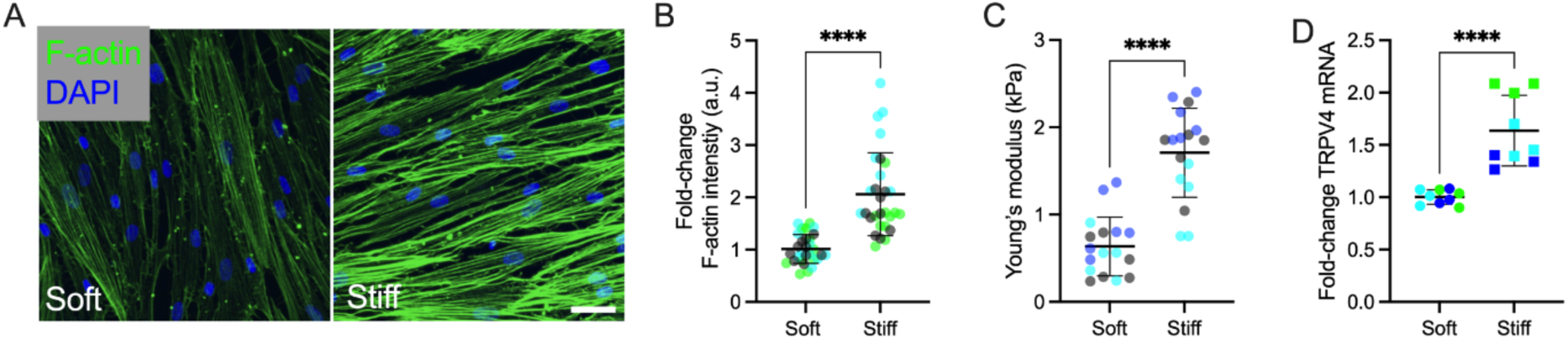
Substrate stiffening increases F-actin and stiffness in nSC cells. **(A)** Representative fluorescence micrographs of labelled F-actin in nSC cells on soft and stiff hydrogel substrates. Scale bars, 20 μm. **(B)** Analysis of F-actin fluorescence intensity (n = 30 images per group from 3 nSC cell strains with three replicates per cell strain). **(C)** Young’s modulus of nSC cells cultured on the soft and stiff substrates measured by AFM (n = 17 cells per group from 3 nSC cell strains). **(D)** Fold-change of mRNA levels for TRPV4 normalized to GAPDH by qRT-PCR (n = 9 replicates from 3 nSC cell strains) in nSC cells. Symbols with the same colors are from the same cell strains. The lines and error bars indicate mean ± SD. Significance was determined by two-way ANOVA using multiple comparisons tests (****p<0.0001).

**Suppl. Fig. 5.**
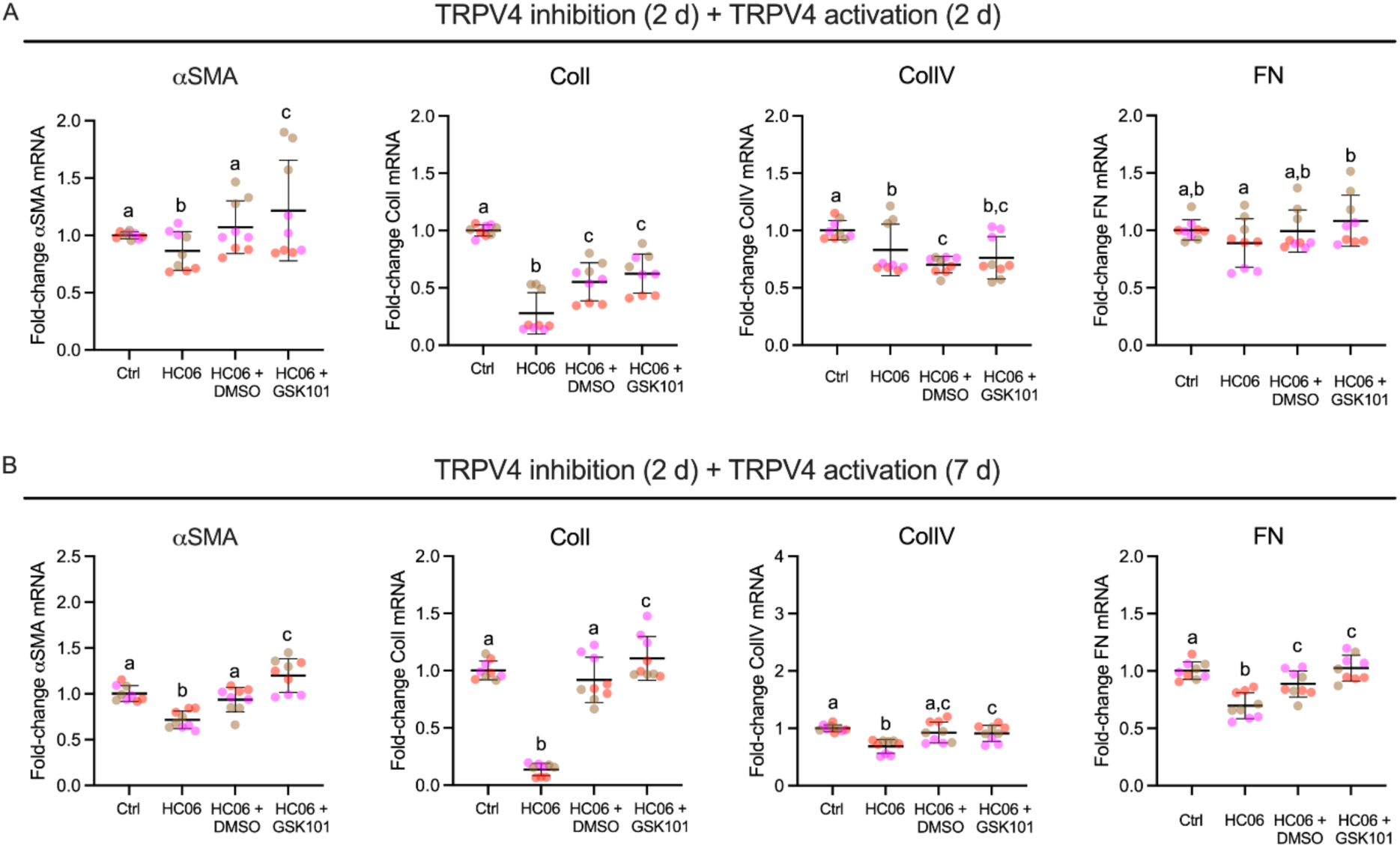
Effects of sustained TRPV4 activity on transcript levels of αSMA and ECM proteins in gSC cells. **(A)** Fold-change of mRNA for αSMA, fibronectin (FN), collagen type I (Col I), and collagen type IV (Col IV) measured by qRT-PCR and normalized to GAPDH in gSC cells under the following conditions: control (DMSO); exposure to the TRPV4 antagonist HC06 (10 μM) for 4 days; exposure to the TRPV4 antagonist HC06 (10 μM) for 2 days, followed by removal of HC06 and exposure to vehicle (DMSO) for 2 days; and exposure to the TRPV4 inhibitor HC06 (10 μM) for 2 days followed by removal of HC06 and exposure to the TRPV4 agonist GSK101 (100 nM) for 2 days (n = 9 replicates from 3 gSC cell strains). **(B)** As in panel (A), except that the duration of exposure to GSK101 or control media was 7 days instead of 2 days. The lines and error bars indicate mean ± SD. Significance was determined by two-way ANOVA using multiple comparisons tests [shared significance indicator letters represent non-significant difference (p > 0.05), distinct letters represent significant difference (p < 0.05)].

**Suppl. Fig. 6.**
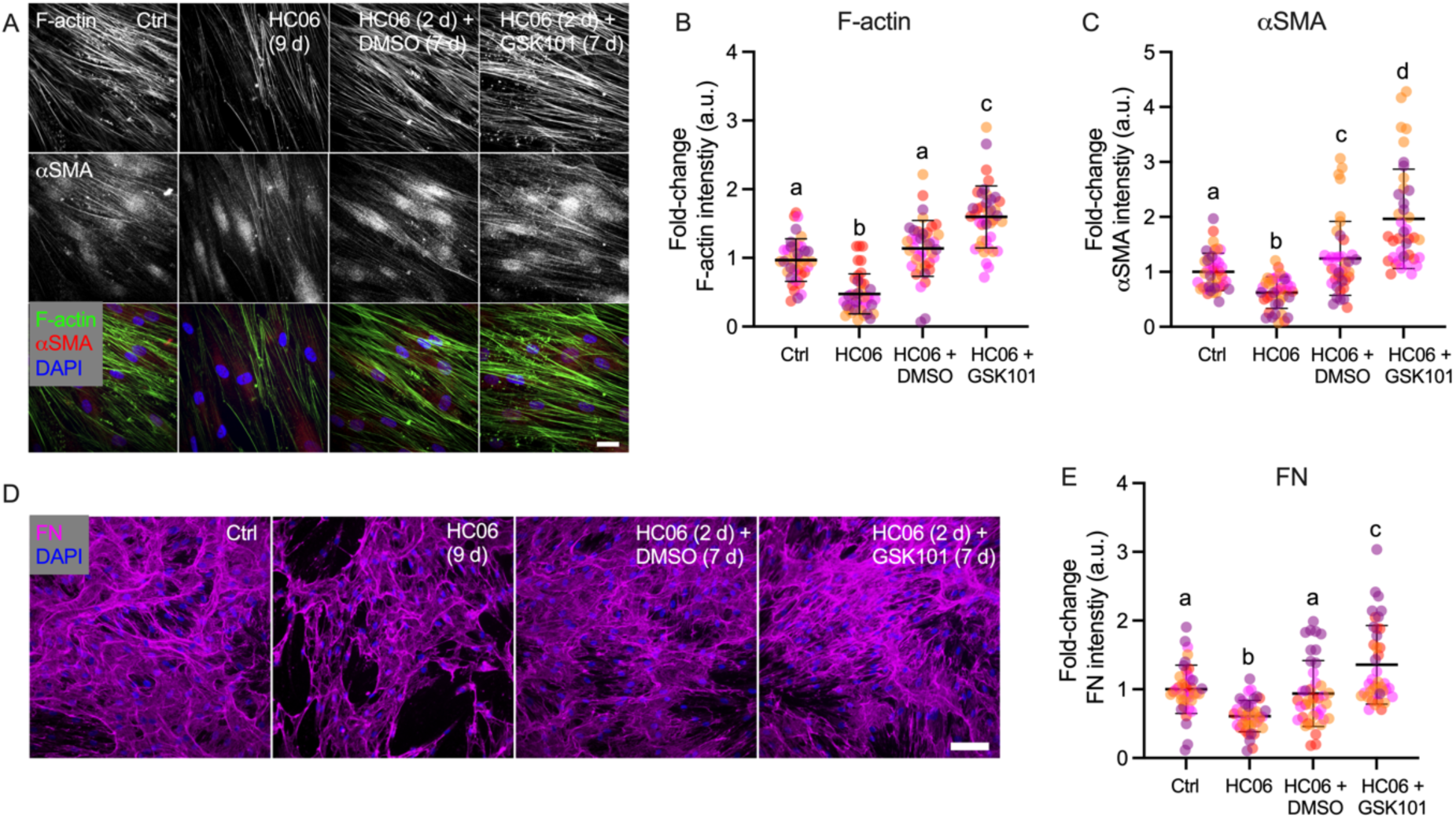
Effects of extended TRPV4 activation on mechanobiological changes in gSC cells. **(A)** Representative fluorescence micrographs of F-actin and αSMA in gSC cells under the following conditions: control (DMSO); exposure to the TRPV4 antagonist HC06 (10 μM) for 9 days; exposure to the TRPV4 antagonist HC06 (10 μM) for 2 days, followed by removal of HC06 and exposure to vehicle (DMSO) for 7 days; and exposure to the TRPV4 inhibitor HC06 (10 μM) for 2 days followed by removal of HC06 and exposure to the TRPV4 agonist GSK101 (100 nM) for 7 days. Scale bar, 20 μm. **(B, C)** Quantification of F-actin and αSMA fluorescent labeling intensities for conditions in panel A (n = 40 images per group from 4 nSC cell strains with three replicates per cell strain). **(D)** Representative fluorescence micrographs of fibronectin (FN) in gSC cells cultured under various conditions as in panel A. Scale bar, 100 μm. **(E)** Quantification of FN fluorescent intensity fr conditions in pabel A (n = 40 images per group from 4 gSC cell strains with three replicates per cell strain. Symbols with different colors represent different cell strains. The lines and error bars indicate Mean ± SD. Significance was determined by two-way ANOVA using multiple comparisons tests [shared significance indicator letters represent non-significant difference (p > 0.05), distinct letters represent significant difference (p < 0.05)].

**Suppl. Fig. 7.**
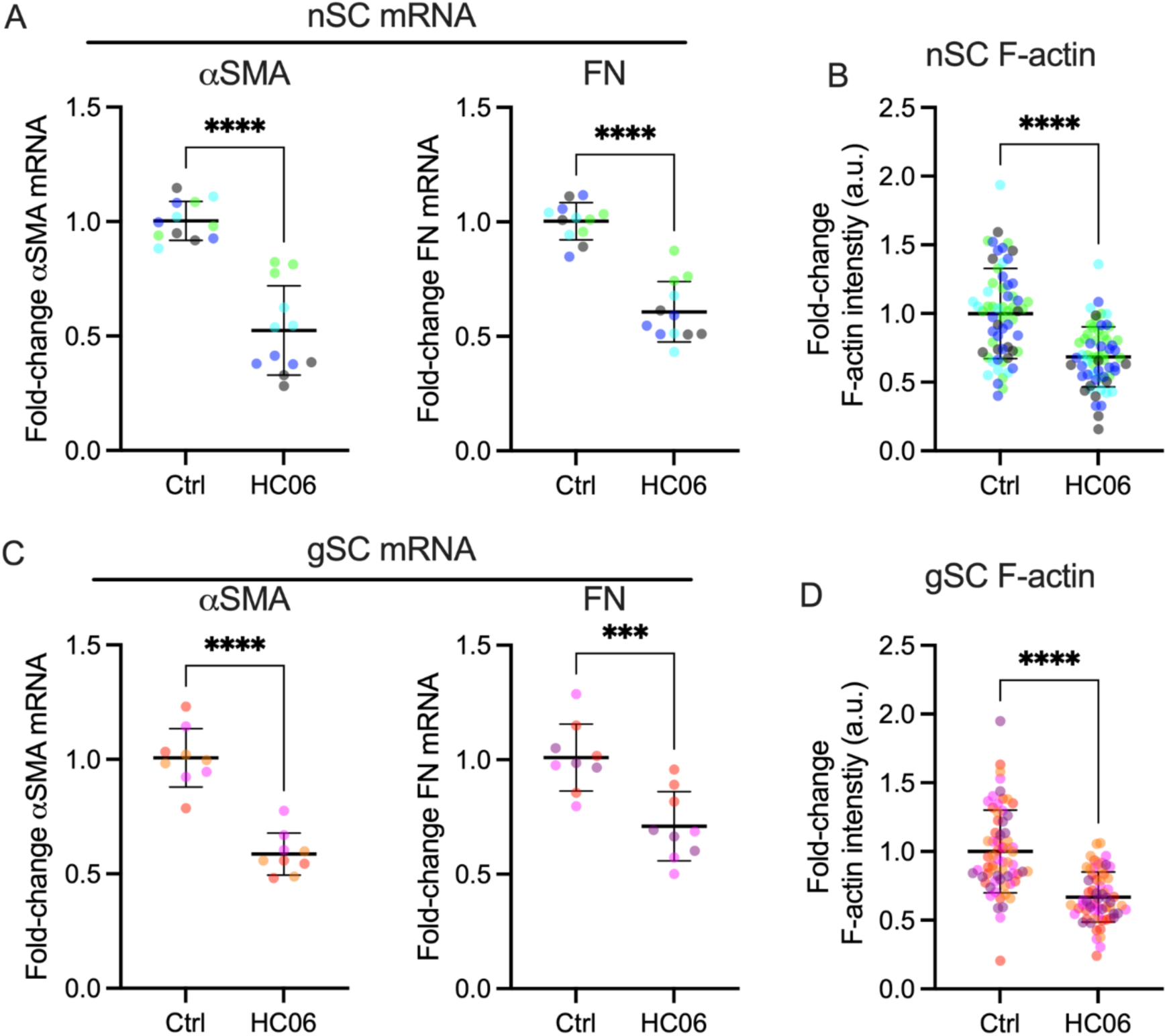
SC cells continue to show reduced actin and fibronectin expression induced by sustained TRPV4 inhibition after trypsinization. **(A, C)** mRNA fold-change of αSMA and fibronectin (FN) normalized to GAPDH by qRT-PCR (n = 9 replicates from 3 nSC/gSC cell strains) in cells treated with or without 10 μM HC06, measured after trypsinization for 2 days. **(B, D)** F-actin intensity (n = 50 images per group from 3 nSC cell strains with three replicates per cell strain; n = 60 images per group from 4 gSC cell strains with three replicates per cell strain), measured after trypsinization and replating on particels. Symbols with different colors represent different cell strains. The lines and error bars indicate Mean ± SD. Significance was determined by two-way ANOVA using multiple comparisons tests (*p<0.05, ****p<0.0001).

**Suppl. Table 1.**
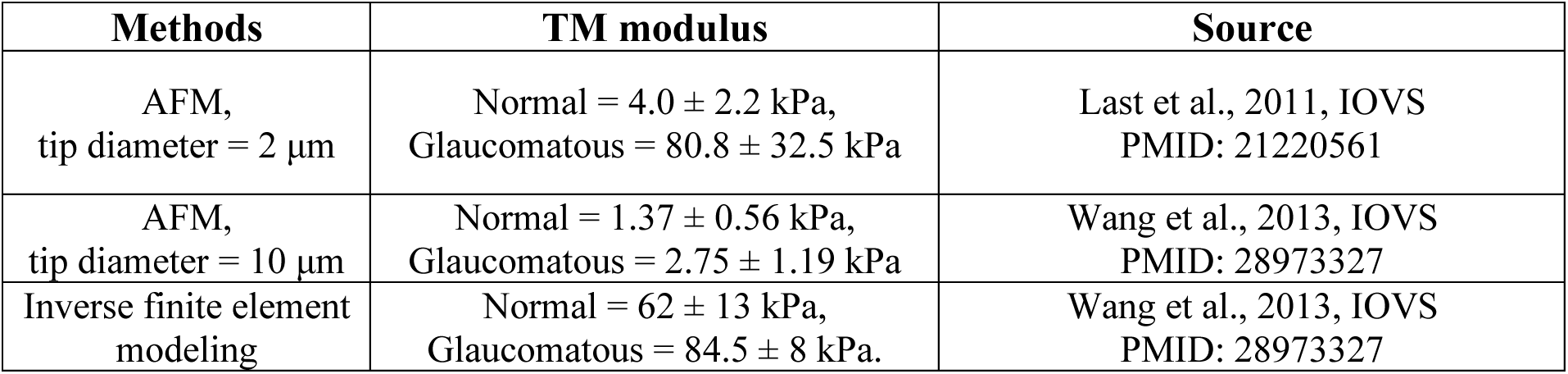
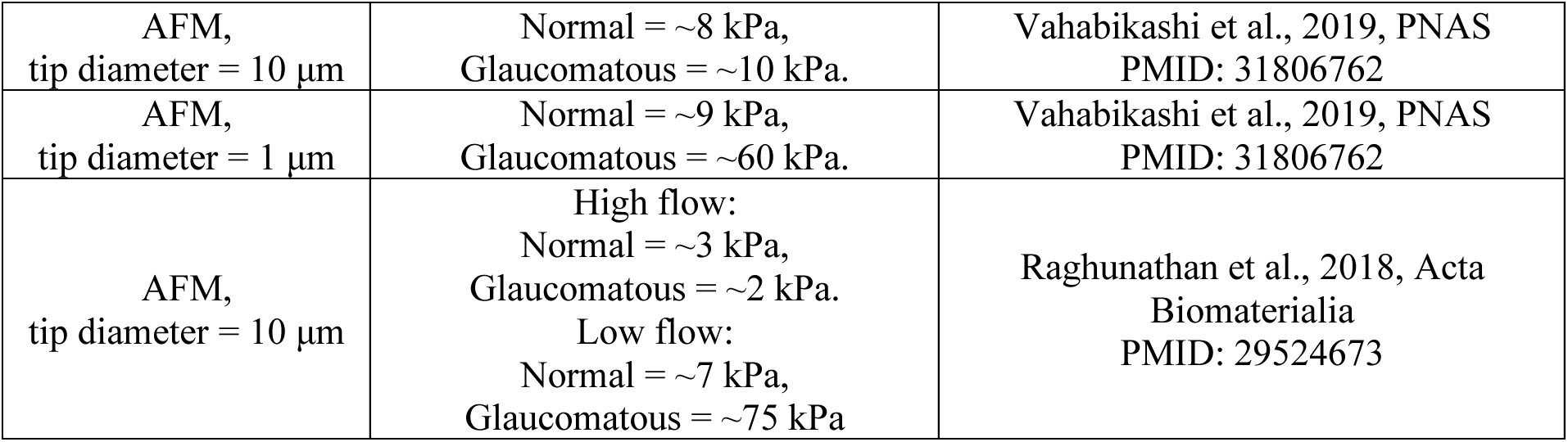
Human trabecular meshwork tissue stiffnesses reported to date.

| Methods | TM modulus | Source |
| --- | --- | --- |
| AFM,<br>tip diameter = 2 μm | Normal = $4.0 \pm 2.2$ kPa,<br>Glaucomatous = $80.8 \pm 32.5$ kPa | Last et al., 2011, IOVS<br>PMID: 21220561 |
| AFM,<br>tip diameter = 10 μm | Normal = $1.37 \pm 0.56$ kPa,<br>Glaucomatous = $2.75 \pm 1.19$ kPa | Wang et al., 2013, IOVS<br>PMID: 28973327 |
| Inverse finite element<br>modeling | Normal = $62 \pm 13$ kPa,<br>Glaucomatous = $84.5 \pm 8$ kPa. | Wang et al., 2013, IOVS<br>PMID: 28973327 |
| AFM,<br>tip diameter = 10 $\mu\text{m}$ | Normal = $\sim 8$ kPa,<br>Glaucomatous = $\sim 10$ kPa. | Vahabikashi et al., 2019, PNAS<br>PMID: 31806762 |
| AFM,<br>tip diameter = 1 $\mu\text{m}$ | Normal = $\sim 9$ kPa,<br>Glaucomatous = $\sim 60$ kPa. | Vahabikashi et al., 2019, PNAS<br>PMID: 31806762 |
| AFM,<br>tip diameter = 10 $\mu\text{m}$ | High flow:<br>Normal = $\sim 3$ kPa,<br>Glaucomatous = $\sim 2$ kPa.<br>Low flow:<br>Normal = $\sim 7$ kPa,<br>Glaucomatous = $\sim 75$ kPa | Raghunathan et al., 2018, Acta<br>Biomaterialia<br>PMID: 29524673 |

